# Human-specific remodeling of an endogenous retrovirus shapes structural diversity at acrocentric nucleolar organizer regions

**DOI:** 10.64898/2026.09.11.751073

**Authors:** Mohammad Waseem, Amit Datta, Hudson O’Neill, Azait Imtiaz, Rafael Contreras-Galindo

## Abstract

Human acrocentric short arms harbor K111, an endogenous retrovirus that became an integral component of nucleolar organizer region (NOR)-associated architecture across all five acrocentric chromosome types. Here we combine complete human and primate genomes, haplotype-resolved population assemblies and parent-offspring trios to reconstruct its evolution and transmission. Predominantly full-length ancestral elements underwent human-specific expansion and remodeling, generating recurrent mosaics and individual-specific multicopy configurations. Population structural variation is concentrated in surrounding satellite landscapes rather than within K111-derived sequences themselves. Trio-resolved genomes reveal direct parental transmission, single-parent remodeling and, crucially, maternal-paternal remodeling in distinct configurations: large reciprocal domains with contrasting parental affinities and de novo sequence exchange within an offspring locus. Raw long reads support both maternal-paternal configurations. K111-derived loci associate with nucleolin-positive compartments. Together, these findings establish K111 as an evolutionarily dynamic component and molecular record of acrocentric NOR architecture, revealing unrecognized remodeling and sequence generation across human generations.

---

The short arms of human acrocentric chromosomes 13, 14, 15, 21 and 22 contain nucleolar organizer regions (NORs), which harbor ribosomal DNA (rDNA) arrays and organize nucleolus formation. Their repetitive composition and extensive sequence similarity historically prevented complete assembly. Telomere-to-telomere and pangenome assemblies have now resolved complex domains containing rDNA, satellite DNA and segmental duplications and revealed extensive homology among nonhomologous acrocentric chromosomes.^1,2^ Population-scale analyses further identified pseudo-homologous regions consistent with recurrent interchromosomal exchange.^3^

Comparative and pedigree studies have extended this view. Complete great-ape genomes reveal lineage-specific reorganization of NOR-bearing chromosomes and their surrounding repeat landscapes.^4^ In humans, conventional allelic recombination is strongly depleted across acrocentric short arms, yet exchange can occur within highly homologous domains. A recent multigenerational study identified de novo ectopic recombination between chromosomes 13 and 21 within an approximately 630-kb region of high sequence identity.^5^ Together, these observations suggest that acrocentric exchange may depend on local sequence architecture and homology rather than uniformly elevated recombination.

K111 is an endogenous retrovirus with an unusual history within this genomic environment. Originally identified as a multicopy sequence related to HERV-K(HML-2), K111 is enriched in centromeric and pericentromeric repeat DNA.^6,7^ Its presence in Pan indicates that the ancestral K111 integration predates the Homo–Pan divergence.^6,7^ These studies also proposed subsequent K111 expansion through recombination and identified related K111/K222 forms.^6,7^ Population studies later revealed variation in K111 copy number and presence.^8^ However, these studies could not resolve the chromosomes carrying these elements, their complete sequence contexts or surrounding repeat architectures.^6–8^ Sequence analysis of human NOR distal-junction regions subsequently placed K111 within centromere repeat (CER)-containing architecture adjacent to rDNA.^9^

Complete acrocentric assemblies now allow K111-family diversification to be examined at chromosome and haplotype resolution and distinguished from remodeling of the larger NOR-associated domains in which these sequences reside. K111 is particularly informative because ancestral K111 and derived K111-family forms can be followed together with their surrounding repeat architecture.

Here we reconstruct K111 evolution across human and great-ape acrocentric chromosomes using comparative genomics, population-scale analyses, parent-offspring trios and cellular imaging. We show that K111 expanded across acrocentric chromosomes in the Homo-Pan ancestral lineage and subsequently underwent extensive human-specific remodeling, producing recurrent sequence mosaics and multicopy configurations across all five human acrocentrics. Trio-resolved analyses further uncover direct parental transmission, single-parent remodeling and maternal-paternal sequence exchange, establishing K111 as a molecular record connecting great-ape evolution, human population diversity and the generation of new acrocentric sequence architectures across generations.

## Results

### K111 expanded across Homo-Pan acrocentrics and diversified in humans

We searched the complete CHM13 T2T assembly using previously characterized K111, K222 and soloLTR sequences and recovered these forms within CER-containing acrocentric regions, together with a truncated soloLTR configuration (Supplementary Data 1). We therefore defined four K111-family structural classes: full-length K111, K222, soloLTR and truncated soloLTR (Fig. 1a). Despite their structural differences, the four classes retain sequence features of the GAATTC target-site duplication, supporting descent from a shared K111 integration. In CHM13, K111 occurs on chromosome 15, K222 on chromosome 13, soloLTRs on chromosomes 14 and 21, and truncated soloLTR on chromosome 22 (Fig. 1b).

**Figure 1.**
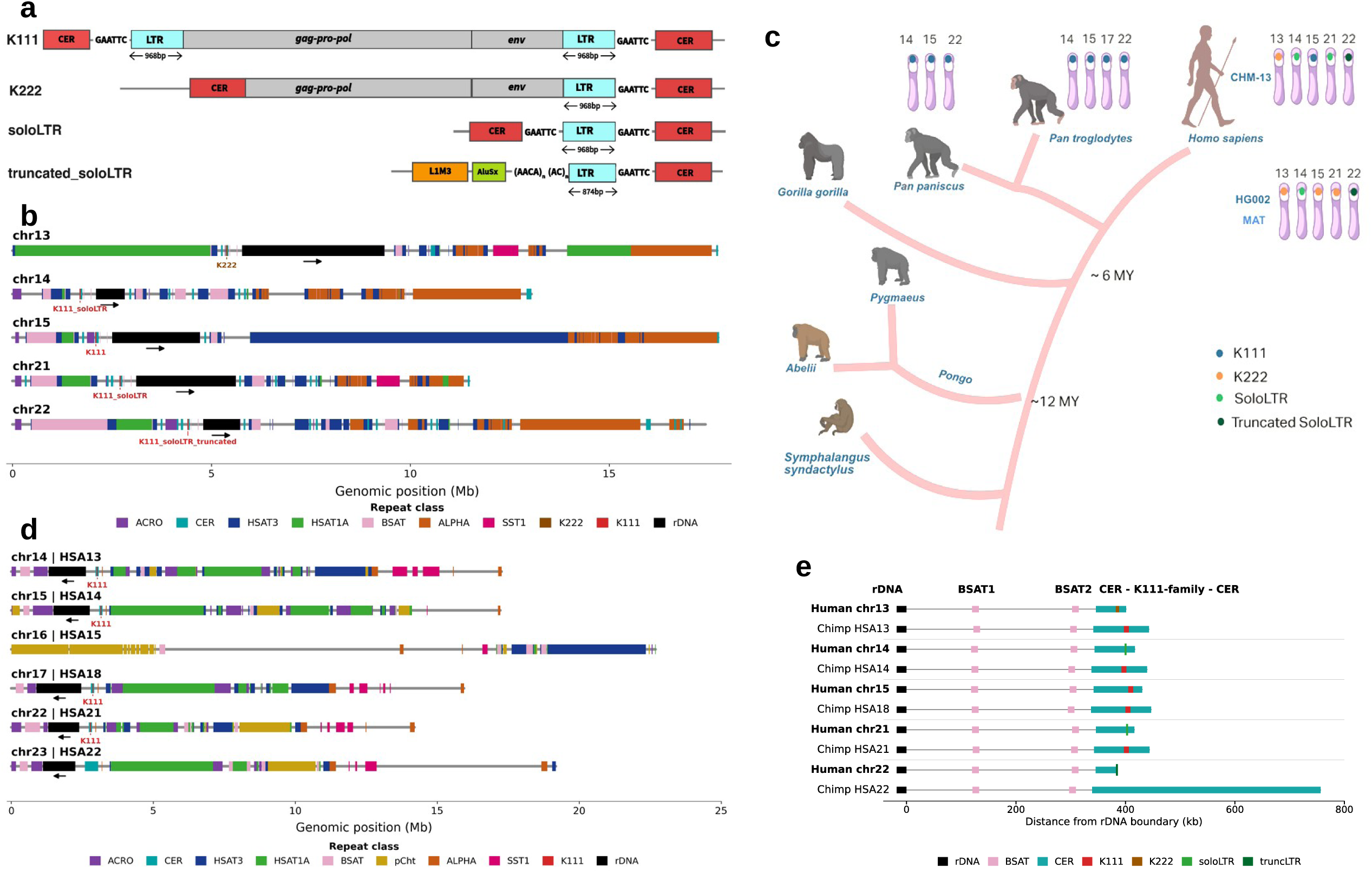
Evolutionary remodeling of K111-family elements at acrocentric nucleolar organizer regions. **a,** Schematic organization of the major K111-family structural classes analyzed in this study: full-length K111, K222, soloLTR and truncated soloLTR. K111 contains 5′ and 3′ LTRs flanking the internal *gag-pro-pol* and *env* regions, whereas K222 lacks the 5′ LTR and soloLTR forms retain only an LTR within the surrounding CER context. The representative truncated soloLTR is shown with adjacent L1M3 and AluSx sequence. Sequence features of the ancestral 6-bp GAATTC target-site duplication are indicated. **b,** Repeat architecture of the five CHM13 human acrocentric short arms (chr13, chr14, chr15, chr21 and chr22), showing the positions of K111-family elements relative to rDNA, CER and other acrocentric repeat classes. Arrows indicate the orientation of the rDNA-containing region. **c,** Evolutionary distribution of K111-family elements across hominoids. Acrocentric chromosomes containing K111-family elements are indicated for chimpanzee (*Pan troglodytes*), bonobo (*Pan paniscus*) and human (*Homo sapiens*); human HG002 MAT illustrates the expanded distribution of K111, K222, soloLTR and truncated soloLTR states across all five acrocentric chromosomes. Approximate divergence times are indicated. **d,** Repeat architecture of chimpanzee chromosome scaffolds homologous to human acrocentrics, showing K111 positions relative to rDNA, CER and other repeat domains. **e,** Comparison of human and chimpanzee NOR-boundary architecture from rDNA through BSAT1, BSAT2 and the CER–K111-family–CER domain. Horizontal distances represent position relative to the rDNA boundary; colors identify CER and the indicated K111-family state. Source data are provided as a Source Data file.

Across available great-ape assemblies, K111 was absent from the examined siamang, orangutan and gorilla genomes but retained in Pan, with four full-length loci in chimpanzee and three in bonobo. By contrast, human acrocentrics contain four structurally distinct K111-family states whose combination and chromosomal distribution also differ between CHM13 and HG002 (Fig. 1c and Supplementary Table 1). These findings place K111 expansion in the Homo–Pan ancestral lineage followed by extensive structural diversification in humans.

Human distal-junction sequence lies telomeric to rDNA, whereas homologous chimpanzee DJ-like sequence and rDNA occur in the opposite orientation.^9^ Consistent with this organization, full-length chimpanzee K111 lies within CER-containing sequence near rDNA (Fig. 1d). Comparative NOR maps showed conservation of the broader rDNA–BSAT–CER organization but substantial divergence within the CER–K111-family module (Fig. 1e and Supplementary Table 2). Four chimpanzee loci retain full-length K111 between CER arrays, whereas corresponding human regions contain K111, K222, soloLTR or truncated soloLTR states accompanied by changes in CER-array length. Chimpanzee HSA22 (HSA, Homo sapiens), for example, lacks K111 and contains an approximately 418-kb rDNA-facing CER block, whereas human chromosome 22 contains an approximately 39-kb CER block adjacent to truncated soloLTR, with the second CER block absent.

NOR architecture did not simply track chromosome homology: chimpanzee chr16 (HSA15) lacks the rDNA-containing NOR domain, whereas chimpanzee chr17 (HSA18) retains an rDNA–BSAT–CER–K111–CER architecture resembling the K111-bearing NOR of human chromosome 15 (Fig. 1d,e). Together, these comparisons support expansion of K111 in the Homo–Pan ancestral lineage within NOR-associated repeat domains, followed by extensive structural and architectural diversification in humans.

### K111-family states diversify across human acrocentric haplotypes

Across 92 haplotype-resolved assemblies from 46 individuals, we identified 455 non-redundant K111-family elements: 73 K111, 169 K222, 194 soloLTR and 19 truncated soloLTR elements (Fig. 2a and Supplementary Table 3). Maternal and paternal structural composition differed in 43 of 46 individuals (93.5%), whereas total K111-family copy number differed in only 21 of 46 and showed no parent-of-origin bias (MAT mean, 5.11; PAT mean, 4.78; paired Wilcoxon test, P = 0.319; Fig. 2a). Thus, haplotype diversity primarily reflects differences in structural composition rather than total element abundance.

**Figure 2.**
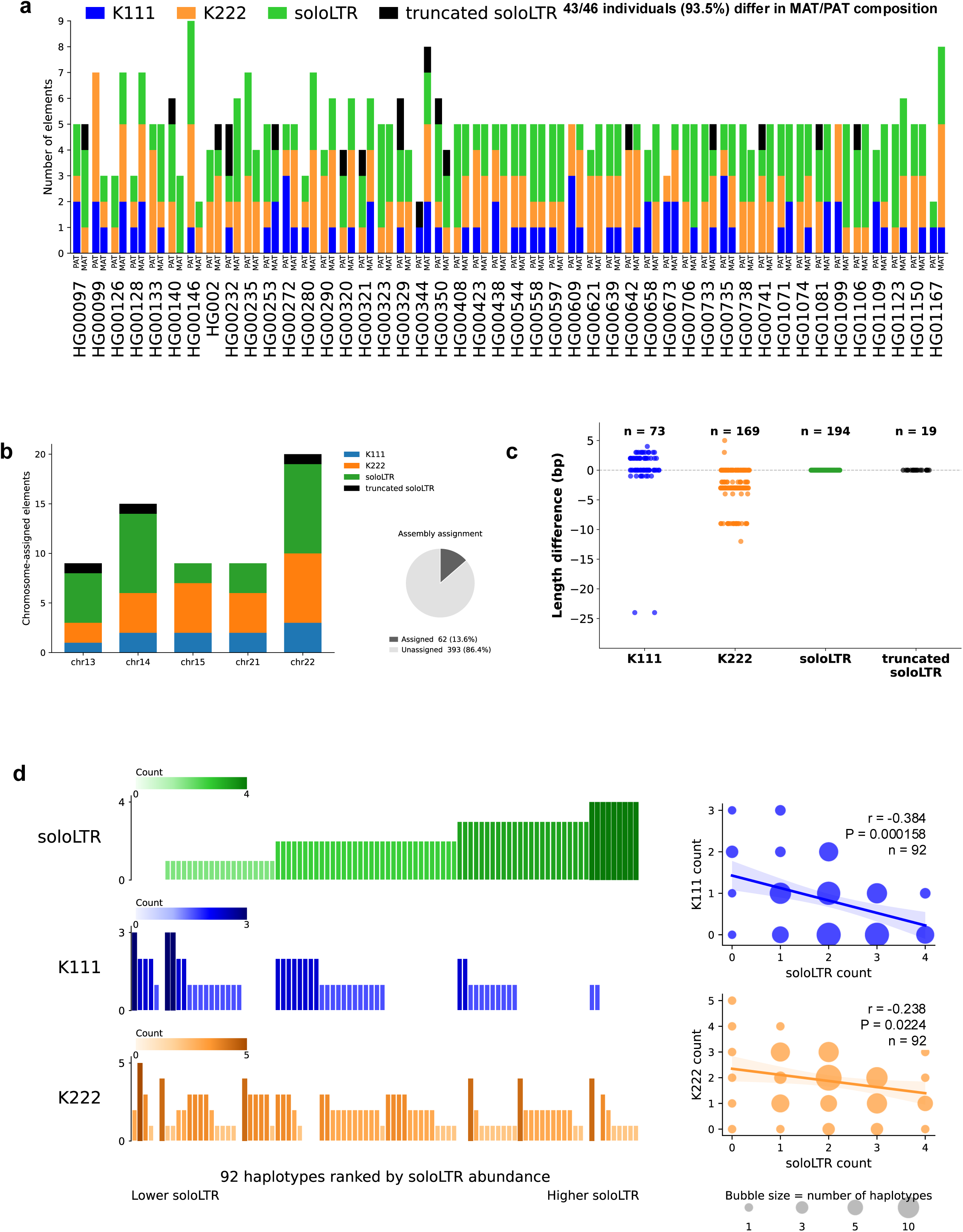
Haplotype-resolved diversity of K111-family elements in the human population. **a,** K111-family composition across 92 maternal (MAT) and paternal (PAT) haplotypes from 46 individuals. Stacked bars show numbers of K111, K222, soloLTR and truncated soloLTR elements per haplotype. MAT/PAT composition differed in 43 of 46 individuals (93.5%). **b,** Chromosome assignment of K111-family elements across chr13, chr14, chr15, chr21 and chr22. Stacked bars show chromosome-assigned elements by structural class; inset summarizes the proportion of elements with and without chromosome-level assignment. **c,** Element-length variation relative to the corresponding reference structure for K111 (*n* = 73), K222 (*n* = 169), soloLTR (*n* = 194) and truncated soloLTR (*n* = 19). Each point represents one element. **d,** Relationship between K111-family classes across the 92 haplotypes. Left, haplotypes ranked by increasing soloLTR abundance, with corresponding K111 and K222 counts shown below. Right, associations between soloLTR abundance and K111 or K222 copy number. K111 abundance was negatively correlated with soloLTR abundance (Pearson *r* = −0.384, *P* = 0.000158), as was K222 abundance (*r* = −0.238, *P* = 0.0224; *n* = 92 haplotypes). Bubble size indicates the number of haplotypes with each combination of element counts. Source data are provided as a Source Data file.

Among 62 elements unambiguously assigned to an acrocentric chromosome, K111-family elements were represented across chromosomes 13, 14, 15, 21 and 22, with no significant difference in the distribution of the four structural classes among chromosomes (χ² = 6.15, d.f. = 12, P = 0.91; Fig. 2b). The remaining 393 elements (86.4%) could not be assigned confidently because of extensive short-arm homology. The chromosome-resolved subset nevertheless demonstrates that K111-family structural states are not restricted to specific acrocentric chromosomes but occur on different acrocentrics across human haplotypes, consistent with exchange among acrocentric short arms^3,5^ and extending beyond the chromosome-specific patterns seen in individual reference genomes.

Structural classes also differed in length variation: K111 ranged from 9,166–9,194 bp and K222 from 6,245–6,262 bp, whereas all observed soloLTR and truncated soloLTR elements were 968 bp and 874 bp, respectively (Fig. 2c). Thus, length variation was restricted to the full-length classes. soloLTR was the most frequent state (194 of 455 elements), and its abundance was inversely correlated with both K111 (Pearson r = −0.384, P = 1.58 × 10^−4^) and K222 abundance (r = −0.238, P = 0.0224; Fig. 2d). These associations are consistent with recurrent remodeling of K111-family architectures but do not alone establish its mechanism.

Finally, K111-family composition did not differ significantly among populations. Four-class composition did not differ among the five represented 1000 Genomes populations (PERMANOVA, P = 0.847), and no individual class remained significantly different after multiple-testing correction (Extended Data Fig. 1). Together, these findings show extensive haplotype-level diversification of K111-family states, with variation driven primarily by structural composition rather than parent-of-origin or population-level effects.

### K111-family sequences show recurrent remodeling and non-random K111/K222 mosaicism

Phylogenetic analysis of 249 K111/K222 internal sequences revealed extensive diversification, with human K111 resolving into nine groups (K111a–i), K222 into 19 groups (K222a–s), and an additional mixed K111/K222 lineage (Extended Data Fig. 2 and Supplementary Data 2). Chimpanzee and bonobo K111 sequences provided evolutionary context for the human lineages.

Full-length comparison distinguished the derived K222 structure from subsequent sequence exchange. Across 168 human K222 elements, the major K222 backbone is collinear with K111 from approximately K111 position 3.06 kb, with minor boundary variation largely attributable to a 3-bp indel (Fig. 3a). Human K111 and all seven Pan K111 sequences retained the corresponding 5′ region, whereas the short sequence preceding the K222–K111 boundary was conserved among K222 elements but could not be assigned to a human or Pan K111 lineage. Thus, the approximately 6.25-kb K222 architecture represents a conserved structural form that likely preceded its subsequent expansion, rather than recurrent independent truncation of K111.

**Figure 3.**
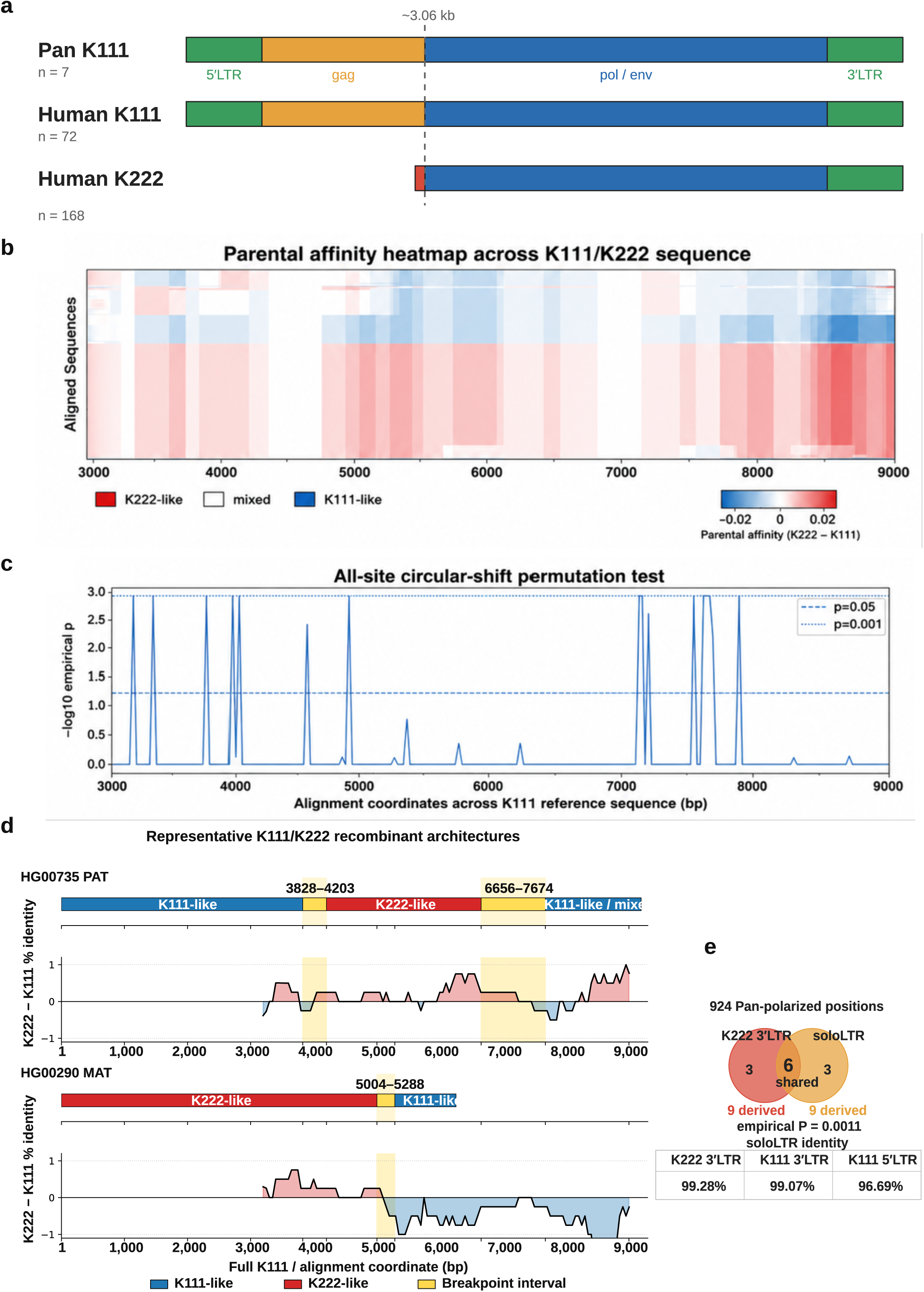
Structural remodeling and recurrent sequence exchange among K111-family elements. **a,** Full-length structural comparison of Pan K111, human K111 and human K222. Human and Pan K111 retain the ∼9.2-kb full-length architecture, whereas 168 evaluable human K222 elements show a conserved ∼6.25-kb structure lacking the 5′ LTR and most of the corresponding K111 5′ region. The principal K222 backbone becomes collinear with K111 at approximately K111 position 3.06 kb. The short K222 sequence preceding this boundary (red) is highly conserved among K222 elements but could not be assigned confidently to either human or Pan K111. **b,** Parental-affinity analysis of 250 aligned K111/K222-derived sequences across K111 positions 3,000–9,000. Colors indicate the sliding-window difference in sequence identity to K222 and K111 consensuses (K222 − K111), with K222-like regions in red and K111-like regions in blue. Windows were 400 bp and required ≥100 informative positions; affinity states were assigned using |Δ identity| ≥0.0025. **c,** Circular-shift permutation analysis of breakpoint localization using 1,000 permutations. Breakpoints were enriched at approximately 3.0–4.0 kb, with a peak at 3,250–3,500 bp containing 58 calls (empirical *P* = 0.001), and at approximately 7.0–7.75 kb, where significant bins contained 51 calls (*P* = 0.001); a smaller enrichment occurred at approximately 4.75–5.0 kb (*P* = 0.018). **d,** SimPlot and RDP5 validation of representative recurrent mosaic architectures. HG00735 PAT contains K111/K222 affinity transitions at 3,828–4,203 and 6,656–7,674 bp, whereas HG00290 MAT contains a transition at 5,004–5,288 bp. SimPlot profiles were calculated using 400-bp windows and 50-bp steps; shaded regions indicate candidate breakpoint intervals. RDP5 analyses independently supported these mosaic architectures across multiple recombination-detection algorithms, with inferred breakpoint intervals overlapping the sliding-window transitions. **e,** Pan-polarized LTR analysis. Among 924 positions at which chimpanzee and bonobo K111 LTR consensuses agreed, K222 3′LTR and soloLTR each contained nine derived states, six of which were shared (one-sided Fisher’s exact test, OR = 608, *P* = 8.23 × 10^−12^; exact circular-shift empirical *P* = 0.00108). The soloLTR consensus was most similar to K222 3′LTR (99.28%), compared with K111 3′LTR (99.07%) and K111 5′LTR (96.69%). Source data are provided as a Source Data file.

Within the approximately 6-kb region shared by K111 and K222, parental-affinity analysis of 250 sequences identified 46 high-confidence mosaics (18.4%) and 287 breakpoint calls (Fig. 3b,c). Breakpoints were non-random, with clustering in the gag/pol region and near the env/3′LTR boundary: circular-shift analysis with 1,000 permutations identified a major peak at 3,250–3,500 bp (58 calls, empirical P = 0.001), enrichment between approximately 7.0 and 7.75 kb (51 calls, P = 0.001), and a smaller peak at 4.75–5.0 kb (P = 0.018; Fig. 3c). Notably, these enriched regions overlap approximate K111/K222 transition regions previously identified in PCR-amplified and sequenced K222/K111 recombinants. These patterns distinguish the conserved structural change defining K222 from recurrent exchange within homologous K111/K222 sequence.

Five elements showed recurrent mosaic architectures: HG00735 PAT and HG01071 PAT showed K111→K222→K111/mixed profiles, whereas HG00290 MAT, HG00344 MAT and HG01074 MAT showed K222→K111 transitions. Representative switches occurred at 3,828–4,203 and 6,656–7,674 bp in HG00735 PAT and 5,004–5,288 bp in HG00290 MAT (Fig. 3d). Complementary RDP5 analyses supported these mosaic architectures across multiple recombination-detection algorithms, with inferred breakpoint intervals overlapping the sliding-window transitions (Fig. 3d).

Phylogenetic analysis of 541 K111-family LTRs further showed diversification across multiple lineages: soloLTRs resolved into eight lineages and K222 3′LTRs into seven, while K111 5′ and 3′LTRs occupied multiple lineages (Extended Data Fig. 3 and Supplementary Data 3). Among 190 correctly oriented full soloLTRs, the consensus was most similar to the K222 3′LTR consensus (962/969 positions; 99.28%), compared with K111 3′LTR (960/969; 99.07%) and K111 5′LTR (936/968; 96.69%). Using 924 Pan-polarized positions, K222 3′LTR and soloLTR each carried nine derived consensus states, six of which were shared (one-sided Fisher’s exact test, OR = 608, P = 8.23 × 10 ¹²; circular-shift empirical P = 0.00108; Fig. 3e), a result that remained significant under a more stringent positional filter. Full-sequence parental modeling independently identified K222 3′LTR as the closest parental class for all 190 soloLTRs, and a K111 5′→3′LTR crossover model did not improve the fit. These results do not establish that soloLTRs arose directly from K222, but support a shared history of derived K111-family LTR remodeling.

### Structural variation is concentrated in repeat domains surrounding K111-family elements

We next examined population structural variation across CHM13 acrocentric short arms using CoLoRSdb v1.0.0. SV density differed markedly among repeat compartments, with enrichment in satellite-rich regions including HSAT1A, HSAT3, BSAT and CER, whereas K111-family sequence was depleted approximately 5.5-fold relative to the remaining analyzed acrocentric sequence (rate ratio = 0.18, FDR q = 5.85 × 10^−6^; Fig. 4a). Thus, structural variation is preferentially concentrated in the repeat domains surrounding K111-family elements rather than within the elements themselves.

**Figure 4.**
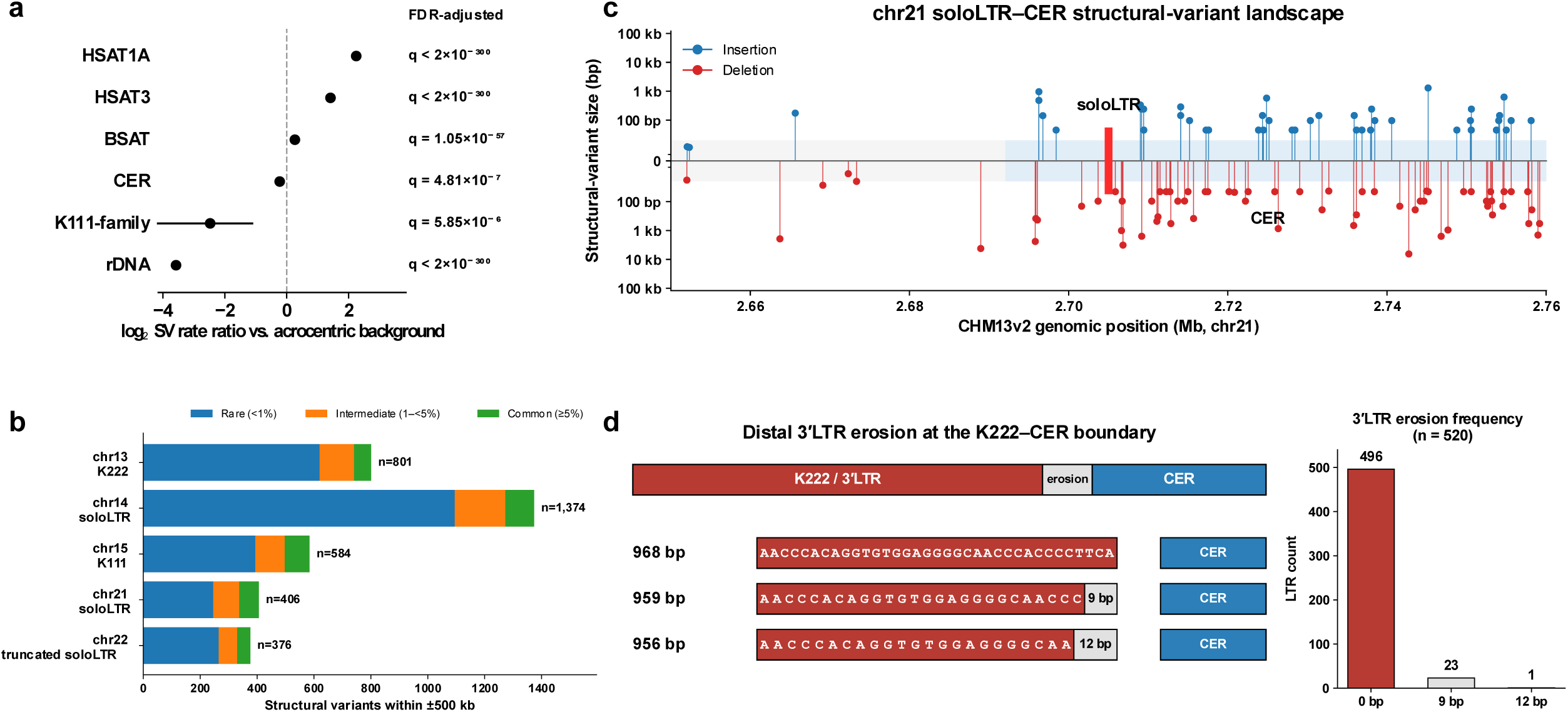
Structural variation is concentrated in repeat domains surrounding K111-family elements. **a**, Structural-variant enrichment across CHM13v2 acrocentric repeat compartments. Rate ratios compare CoLoRSdb v1.0.0 SV density within each repeat class with the remaining analyzed acrocentric sequence; points indicate rate ratios and horizontal lines 95% confidence intervals. Two-sided conditional binomial rate comparisons were performed from SV counts and analyzed sequence lengths, with Benjamini–Hochberg correction across six repeat classes; FDR-adjusted q values are shown. **b,** Structural-variant burden and allele-frequency composition within ±500 kb of the five CHM13 K111-family loci. Rare variants have AF <1%, intermediate variants AF 1–<5% and common variants AF ≥5%. **c,** Structural-variant landscape across a 110-kb window centered on the chr21 soloLTR. Insertions and deletions are plotted above and below the baseline, respectively; CenSat-annotated CER sequence is shown in light blue and the 968-bp soloLTR in red. **d,** Distal 3′ LTR erosion at K111-family–CER junctions, showing reference-length, 9-bp and nested 12-bp erosion states. Among 520 non-truncated LTRs, 23 carried the 9-bp erosion and one carried the 12-bp erosion; together these variants occurred in 18 of 46 individuals (39%). CoLoRSdb v1.0.0 contained 1,381 samples with data. Source data are provided as a Source Data file.

SV burden and allele-frequency composition also varied among the five CHM13 K111-family loci across ±500-kb windows, with variants spanning rare (AF <1%), intermediate (AF 1–<5%) and common (AF ≥5%) frequency classes (Fig. 4b). Examination of 110-kb windows around each locus identified 90, 138, 168, 110 and 26 SVs around the chromosome 13 K222, chromosome 14 soloLTR, chromosome 15 K111, chromosome 21 soloLTR and chromosome 22 truncated soloLTR loci, respectively, whereas only 3, 0, 1, 0 and 0 variant start sites occurred within the corresponding K111-family elements (Fig. 4c and Extended Data Fig. 4).

Quantitative comparison with the local repeat environment confirmed this pattern: K111-family cores showed a 6.5-fold lower SV density than CenSat-annotated CER within 50 kb of the elements (rate ratio = 0.155, 95% CI = 0.042–0.399, P = 5.66 × 10□□; Extended Data Fig. 5). By contrast, SV density in CER within 0–5 kb of K111-family boundaries did not differ from CER 5–50 kb away (rate ratio = 0.90, P = 0.49), arguing against preferential structural instability at the retroviral boundaries.

Localized remodeling of K111-family junctions was also evident as distal 3′LTR erosion occurring in stereotyped nested forms. Among 520 non-truncated LTRs, 23 carried a 9-bp distal erosion (4.4%) and one carried a nested 12-bp erosion (0.2%); together these variants were detected in 18 of 46 individuals (39%) (Fig. 4d). These recurrent junctional changes, together with the population SV landscape, distinguish relatively constrained K111-family sequence from the substantially more variable repeat architecture surrounding it.

### Recurrent K111-family loci occur within remodeled NOR-associated domains

Six haplotype-resolved scaffolds contained two K111-family loci (Fig. 5a and Extended Data Fig. 6a). Five pairs were separated by 1.46–3.30 Mb, whereas the two K222 elements on HG00235 PAT were only 24.3 kb apart. CER abundance differed between paired loci on five of six haplotypes, and the 12 individual elements varied substantially in both CER abundance and sidedness within ±100 kb (Extended Data Fig. 6b,c). Thus, recurrent K111-family loci occur within CER-associated environments whose local repeat architecture is not fixed.

**Figure 5.**
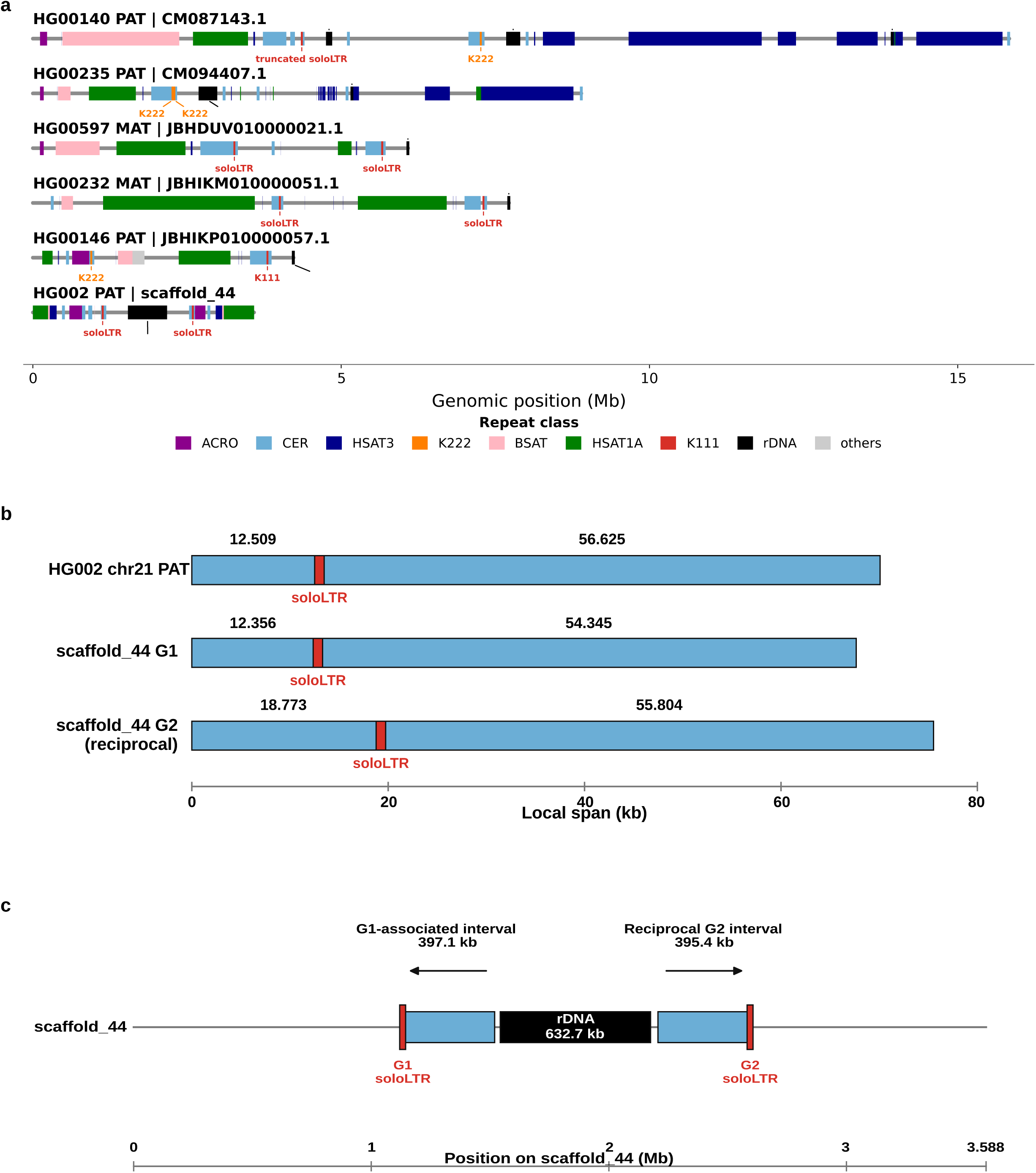
Recurrent K111-family loci mark extended, remodeled NOR-associated repeat domains. **a,** Repeat architecture of six haplotype-resolved scaffolds containing two K111-family loci: HG00140 PAT (CM087143.1), HG00235 PAT (CM094407.1), HG00597 MAT (JBHDUV010000021.1), HG00232 MAT (JBHIKM010000051.1), HG00146 PAT (JBHIKP010000057.1) and HG002 PAT (scaffold_44). Major acrocentric repeat classes, rDNA and K111-family elements are indicated. **b,** Local CER–soloLTR architecture of the HG002 paternal chromosome 21 locus and scaffold_44 G1 and reciprocal G2. Values above CER segments indicate their lengths. G1 closely reproduces the chromosome 21 architecture, whereas G2 contains expansion of the orientation-normalized short CER block. **c,** Extended HG002 paternal scaffold_44 architecture. A 397.1-kb G1-associated interval and a 395.4-kb reciprocal G2 interval occur in opposite orientations on either side of a 632.7-kb rDNA array. The intervals showed 76.1% exact 31-mer correspondence (1,000,000 scaffold-aware matched-length permutations; empirical P = 0.001835), and the G1 and reverse-complemented G2 soloLTRs were 99.79% identical. Arrows indicate interval orientation. Source data are provided as a Source Data file.

The HG002 paternal scaffold_44 provided a resolved example. The paternal chromosome 21 soloLTR was flanked by 12.509-kb and 56.625-kb CER blocks, closely reproduced at scaffold_44 G1 (12.356-kb CER–soloLTR–54.345-kb CER; Fig. 5b). After orientation normalization, the reciprocal G2 region retained a similar long CER block (55.804 kb) but an expanded short block (18.773 kb), 6.264 kb larger than its chromosome 21 counterpart. These relationships identify a recurrent CER–soloLTR architecture accompanied by asymmetric remodeling of its flanking CER blocks.

Similarity between G1 and G2 extended far beyond the soloLTR. A 397.1-kb G1-associated interval and a 395.4-kb reciprocal G2 interval occurred in opposite orientations on either side of a 632.7-kb rDNA array (Fig. 5c). The intervals showed 76.1% exact 31-mer correspondence, defined as the fraction of 31-bp sequences in one interval with an exact sequence match in the reciprocal interval, significantly exceeding the scaffold-aware matched-length null expectation (1,000,000 permutations; empirical P = 0.001835), and the G1 and reverse-complemented G2 soloLTRs were 99.79% identical. Despite this extended reciprocal homology, the intervals were not mirror images in repeat composition, indicating local remodeling within the homologous domains. Together, these findings show that recurrent K111-family loci can mark hundreds-of-kilobase homologous domains embedded within broader, structurally remodeled NOR-associated architecture rather than invariant duplicated K111–CER units.

### Trio analysis identifies transmitted and remodeled K111-family architectures

To distinguish parental transmission from local architectural divergence, we evaluated six parent–offspring trios and performed detailed trio-resolved analysis in HG002 and HG00733, the two trios with sufficient long-read coverage (>20×) and sufficiently complete parental K111-family haplotypes for reliable offspring-to-parent comparisons. Sequence and repeat-architecture similarity were integrated into a transmission score, whereas remodeling was measured by comparing each offspring locus with the range of architectural variation observed among loci from the corresponding phased parent (Fig. 6a,b).

**Figure 6.**
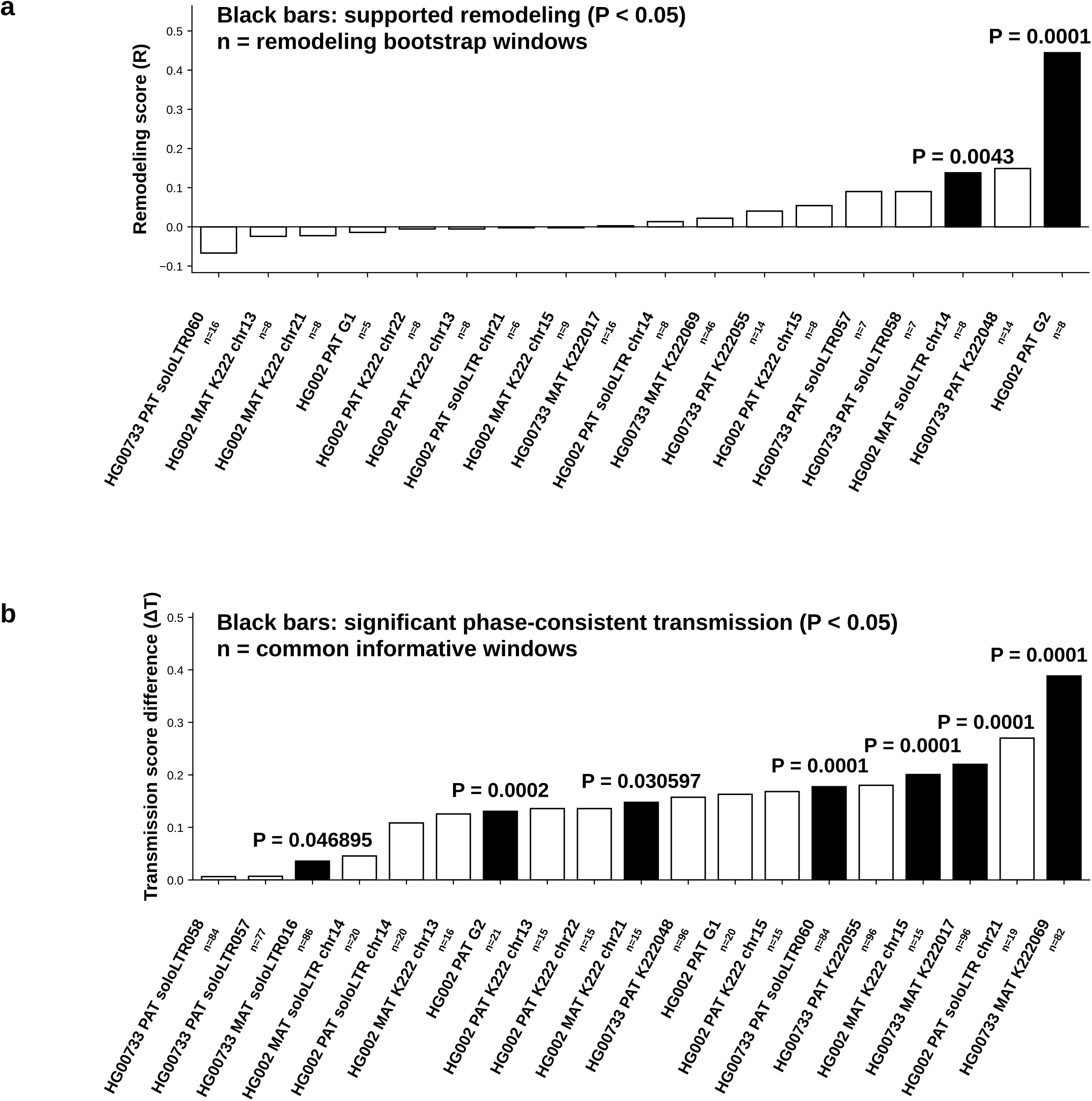
Trio analysis identifies transmitted and remodeled K111-family architectures. **a,** Structural remodeling scores (R) for evaluable K111-family loci in the HG002 and HG00733 trios. Remodeling measures excess offspring-to-parent architectural divergence relative to variation among representative loci from the phased parent. Black bars denote loci with R > 0 and bootstrap P < 0.05. HG002 maternal chromosome 14 soloLTR (R = 0.138, P = 0.0043) and paternal scaffold_44 G2 (R = 0.445, P = 0.0001) showed supported remodeling, whereas scaffold_44 G1 did not (R = −0.014, P = 0.378). **b,** Difference in composite transmission score (ΔT) between the best-supported and alternative parental architectures. Black bars indicate significant parental discrimination for which the best-supported match was consistent with phased parental origin (bootstrap P < 0.05). Significant cross-parent similarities are shown as open bars and were not classified as transmission. scaffold_44 G2 showed a phase-consistent paternal match to HG003 G3 (P = 0.0002), whereas G1 showed a cross-parent similarity to an HG004 maternal locus. G2 was the only tested locus with both phase-consistent transmission support and supported remodeling. n denotes the number of informative windows. Source data are provided as a Source Data file.

Most evaluable offspring loci showed no significant excess remodeling (Fig. 6a). In HG002, the maternal chromosome 14 soloLTR showed significant remodeling (R = 0.138, bootstrap P = 0.0043). A stronger signal occurred at scaffold_44 G2, which had the largest remodeling score among tested loci (R = 0.445, P = 0.0001), whereas neighboring G1 showed no remodeling excess (R = −0.014, P = 0.378). Thus, the two homologous scaffold_44 domains differed markedly in their relationship to parental repeat architectures.

Transmission analysis identified parental similarities consistent with phased haplotype origin (Fig. 6b). K222 and soloLTR loci showed significant phase-consistent support. At scaffold_44, G2 was most similar to paternal HG003 G3 and showed significant phase-consistent support (bootstrap P = 0.0002), whereas G1 favored an HG004 maternal locus and was therefore retained as a cross-parent similarity rather than classified as paternal transmission. G2 was the only tested locus showing both significant phase-consistent transmission support and significant remodeling.

Local parental-affinity analysis further distinguished the reciprocal scaffold_44 domains (Extended Data Fig. 7). Among non-ambiguous 250-bp windows, G1 contained 24 maternal-like and four paternal-like windows, whereas G2 contained one maternal-like and 49 paternal-like windows (two-sided Fisher’s exact test, P = 6.01 × 10 ¹ ). Independent ONT data supported the assembled G2 organization, with 44 R9 and 70 R10 molecules spanning the CER–bridge–ACRO interval. Among these, four R9 and six R10 molecules traversed the complete CER–soloLTR–CER–intervening sequence–ACRO path (Extended Data Fig. 7). Because the intervening rDNA array prevents nucleotide-resolution transition mapping, these data establish contrasting maternal-and paternal-like sequence ancestry across independently validated reciprocal domains but do not localize an exchange boundary or establish its molecular mechanism.

Together, trio-wide analysis distinguishes largely concordant parental transmission from locus-specific architectural remodeling and identifies scaffold_44 as a domain containing sharply contrasting parental sequence affinities between neighboring loci.

### Raw molecules validate de novo maternal–paternal sequence exchange within an HG00733 acrocentric locus

Across the 475,435-bp HG00733 scaffold-048 interval, assembly-based parental-affinity mapping identified two significant reciprocal ancestry transitions within the K222-containing repeat domain (Fig. 7a,b). The first occurred at position 229,125, changing from paternal-like to maternal-like sequence ancestry (circular-shift permutation P = 0.000600). The second occurred at position 348,375, changing from maternal-like to paternal-like ancestry (P = 0.019398), thereby defining the distal boundary of a maternal-like tract overlapping ACRO sequence. This paternal-like→maternal-like→paternal-like pattern is consistent with a single recombinant tract but does not exclude two independent exchange events.

**Figure 7.**
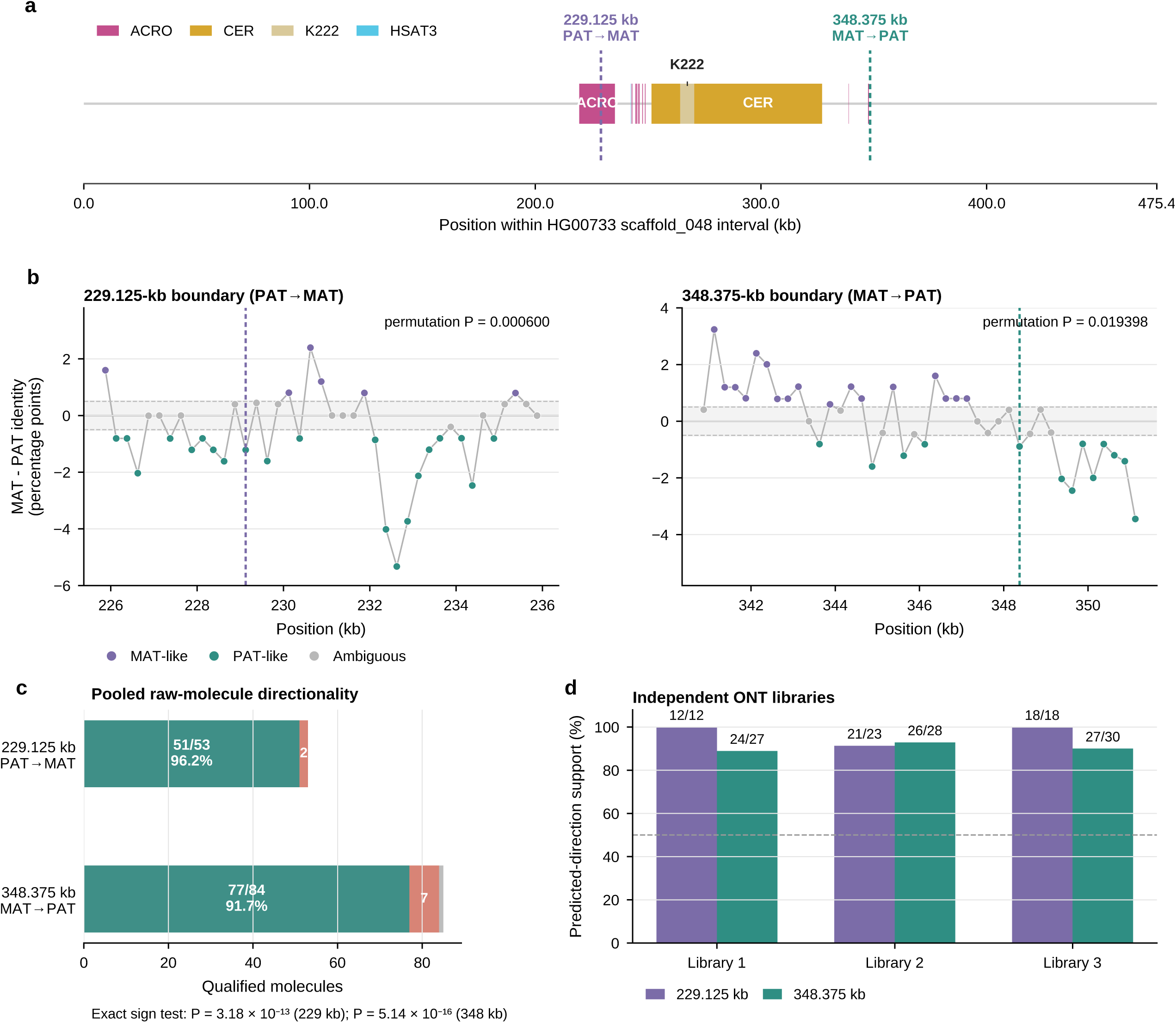
Raw molecules validate de novo maternal–paternal sequence exchange within an HG00733 acrocentric locus. **a,** Repeat architecture across the 475,435-bp HG00733 paternal scaffold 048 interval, showing the K222-associated CER and ACRO domains and the supported ancestry transitions at positions 229,125 and 348,375. **b,** Assembly-based parental-affinity profiles surrounding the two boundaries. The transition at 229,125 changes from paternal-like to maternal-like affinity (circular-shift permutation P = 0.000600), whereas the transition at 348,375 changes from maternal-like to paternal-like affinity (P = 0.019398). **c,** Raw-molecule directionality. At 229,125, 51 of 53 molecules supported the predicted direction and two supported the opposite direction (exact two-sided sign test, P = 3.18 × 10^−13^). At 348,375, 77 of 84 non-tied molecules supported the predicted direction and seven supported the opposite direction (P = 5.14 × 10^−16^); one tied molecule was excluded. **d,** Predicted-direction support stratified across three independent ONT libraries. Source data are provided as a Source Data file.

Raw ONT molecules independently supported the 229,125 transition (Fig. 7c and Extended Data Fig. 8a). Among 53 molecules containing at least four parental-informative sites on each side of the transition, 51 shifted in the predicted paternal-like to maternal-like direction and two in the opposite direction (96.2%; exact two-sided sign test, P = 3.18 × 10□¹³). Support was reproduced across all three independent libraries (12/12, 21/23 and 18/18 molecules).

The reciprocal 348,375 transition was similarly supported (Fig. 7c,d and Extended Data Fig. 8b). After excluding one tie, 77 of 84 molecules shifted in the predicted maternal-like to paternal-like direction and seven in the opposite direction (91.7%; exact two-sided sign test, P = 5.14 × 10□¹□), with concordant support across all three libraries (24/27, 26/28 and 27/30 molecules).

Together, assembly-level permutation tests and molecule-level analyses support two reciprocal ancestry transitions within the HG00733 acrocentric interval. Their recovery on individual offspring molecules across three independent ONT libraries argues against mapping or haplotype-assembly artifacts. The paternal-like→maternal-like→paternal-like pattern is consistent with a single recombinant tract, although two independent exchange events cannot be excluded. These data provide direct molecule-level evidence for de novo maternal–paternal acrocentric sequence exchange during transmission.

### K111-derived sequences associate with nucleolar compartments

Because K111-family elements occupy NOR-bearing acrocentric short arms, we asked whether structurally distinct K111-derived loci share a nucleolar-associated nuclear environment. IF–FISH in BJ-5Ta cells using K111-, K222-and soloLTR-discriminating probes with nucleolin immunofluorescence showed signals from all three classes associated with nucleolin-positive domains (Fig. 8).

**Figure 8.**
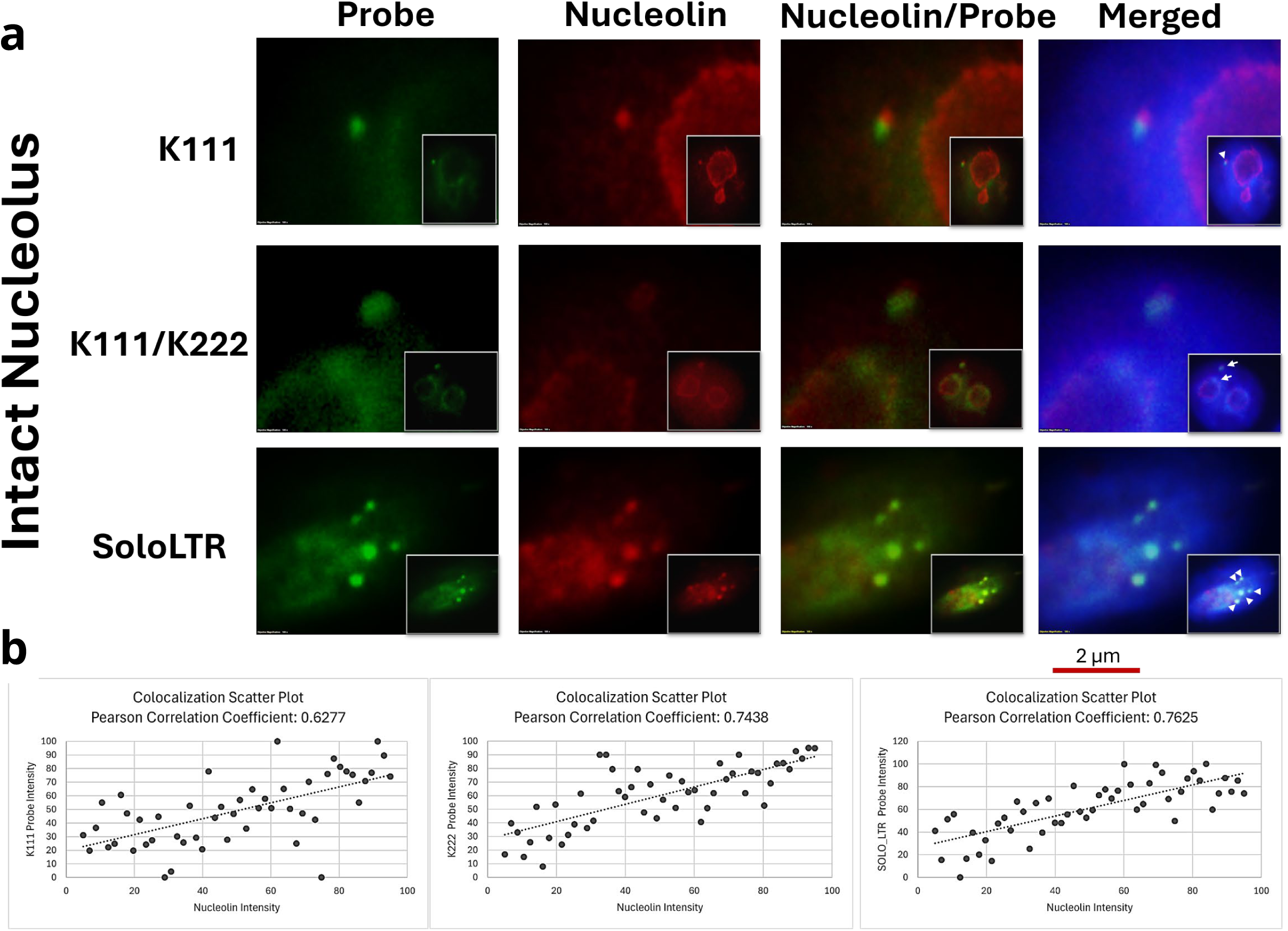
K111-derived sequences associate with nucleolar compartments. **a,** Representative IF–FISH images of K111 (top), K222 (middle) and soloLTR (bottom) in BJ-5Ta cells. K111-family FISH signals are shown in green, nucleolin in red and DNA in blue (DAPI); merged images show their spatial relationship. Insets show representative nuclei and arrowheads indicate K111-family signals associated with nucleolin-positive domains. Scale bar, 2 µm. **b,** Scatter plots show paired probe and nucleolin fluorescence intensities from 50 probe-associated ROIs per sequence class. Pearson correlation coefficients were r = 0.6277 for K111, r = 0.7438 for K222 and r = 0.7625 for soloLTR. Data were obtained from three independent experiments. Source data are provided as a Source Data file.

Across 50 probe-associated regions of interest (ROIs) per sequence class, probe and nucleolin fluorescence intensities were positively correlated for K111 (Pearson r = 0.6277), K222 (r = 0.7438) and soloLTR (r = 0.7625) (Fig. 8). Thus, despite their distinct sequence architectures and evolutionary histories, all three K111-family states show association with nucleolin-positive compartments. Measurements were obtained across three independent experiments.

K111, K222 and soloLTR signals also remained associated with redistributed nucleolin-positive domains in cells exhibiting disrupted nuclear and nucleolar organization following cell detachment and cytospin preparation (Extended Data Fig. 9). This qualitative observation provides orthogonal support for their nucleolar association but was not interpreted as a mechanistic perturbation.

Together, these findings place structurally diverse K111-derived loci within nucleolar-associated nuclear environments, providing a spatial context compatible with interactions among NOR-associated acrocentric sequences without establishing that nucleolar proximity drives sequence exchange.

## Discussion

Complete human and ape assemblies have transformed acrocentric short arms from inaccessible heterochromatin into sequence-resolved regions, revealing homology among heterologous acrocentrics, pseudo-homologous regions consistent with interchromosomal exchange and rapid turnover of NOR-bearing chromosomes.^1–4^ Here, K111 provides a molecular record of these processes. Rather than simple amplification of an ancestral insertion, K111-family evolution involved structural diversification and recurrent sequence exchange within extensively remodeled NOR-associated repeat domains.

Comparative analysis places this diversification in evolutionary context. K111 was not detected in the siamang, orangutan or gorilla assemblies examined, whereas full-length K111 is retained at multiple Pan NOR-associated loci and human acrocentrics contain K111, K222, soloLTR and truncated soloLTR states. This distribution places expansion across acrocentric chromosomes in the Homo-Pan ancestral lineage, followed by extensive human-specific remodeling. NOR architecture does not track simple chromosome orthology across humans and Pan, consistent with evolutionary turnover of ape NOR-bearing chromosomes.^4^ Greater haplotype-resolved sampling in humans means that additional diversity may exist within ape populations. In humans, K111-family states occur across all five acrocentrics, and several haplotypes contain multiple loci in distinct NOR-associated configurations, demonstrating diversification of both sequence composition and large-scale architecture. Their broad distribution and absence of significant population-level differences are consistent with historical exchange among homologous acrocentric domains.

Sequence diversification separates the origin of K111-family structural states from subsequent exchange among them. K222 retains a conserved derived architecture, arguing against recurrent independent truncation of K111, whereas mosaic K111/K222 sequences show non-random transitions concentrated in specific regions. LTR evolution further supports derived K111-family remodeling. Earlier studies proposed recombination during K111 expansion but lacked chromosome-and haplotype-resolved context.^6,7^ Our data distinguish formation of the conserved K222 architecture from recurrent exchange among established K111-derived lineages and place both within resolved acrocentric repeat domains.

These domains are heterogeneous. Structural variants were enriched in surrounding satellite compartments, whereas K111-family cores were comparatively depleted and showed no excess instability at their boundaries. K111-derived sequences therefore reside within dynamic satellite environments rather than behaving as local variation hotspots. Recurrent K111-family loci also marked homology extending beyond the retroviral sequence. On HG002 scaffold_44, reciprocal domains spanning hundreds of kilobases share extensive sequence correspondence but differ in repeat composition, arguing against invariant duplication of small K111-CER units. Such extended homology is relevant because pseudo-homologous regions and pedigree analyses identify long, highly similar acrocentric domains as substrates for interchromosomal exchange.^3,5^ The trio analyses extend this evolutionary picture into contemporary transmission. Most evaluable loci retained parental architectures, while others showed remodeling within single-parent backgrounds. Interestingly, additional loci revealed maternal-paternal remodeling in distinct configurations. HG002 scaffold_44 combined contrasting parental affinities with locus-specific remodeling across large reciprocal domains, whereas HG00733 contained two reciprocal maternal-paternal ancestry transitions within an individual offspring locus. Independent raw long reads support both the scaffold_44 architecture and HG00733 transitions. The paternal-like to maternal-like to paternal-like pattern in HG00733 is consistent with a single recombinant tract, although two independent exchange events cannot be excluded. Together, these observations reveal direct transmission, single-parent remodeling and maternal-paternal remodeling, including sequence-and molecule-level evidence for de novo exchange during transmission.

These findings complement evidence that conventional allelic recombination is strongly depleted across acrocentric short arms despite occasional ectopic exchange within highly homologous domains.^5^ Lin et al. identified a chr13-chr21 ectopic recombination event within an approximately 630-kb region of high sequence identity, whereas population analyses identified pseudo-homologous regions consistent with recurrent heterologous exchange.^3,5^ These observations argue against uniformly recombination-prone acrocentric short arms. Instead, exchange appears to depend on local sequence identity and repeat architecture. The extended homologous domains and distinct maternal-paternal configurations identified here support this architecture-dependent model.

Nuclear organization provides a plausible spatial context. NORs from different acrocentrics converge within nucleoli, bringing heterologous chromosomes into proximity.^3,9–12^ K111, K222 and soloLTR signals associated with nucleolin-positive compartments despite their different architectures and histories. This does not establish that nucleolar proximity causes recombination, but it places K111-family loci where multiple NOR-bearing chromosomes converge. The possible relationship among spatial organization, repeat architecture and acrocentric rearrangements^13^ is considered in Supplementary Discussion 1.

Together, our findings support a model in which acrocentric evolution reflects extensive sequence homology, dynamic satellite architecture and spatial convergence of NOR-bearing chromosomes. K111 is not simply an endogenous retrovirus that expanded across acrocentrics in the Homo-Pan ancestral lineage. Its human-specific diversification, population-level multicopy architectures and remodeling during transmission provide a sequence-resolved record of the processes shaping human NOR-associated short arms.

The number of resolved parent-offspring acrocentric haplotypes remains limited, and larger multigenerational datasets are needed to determine the frequency, chromosome specificity and mechanisms of remodeling and exchange. Our data do not establish that K111 initiates recombination, that HG00733 arose through a particular pathway or that nucleolar association drives exchange. For scaffold_44, the repetitive rDNA-containing interval prevents precise localization of a potential exchange boundary. Future pedigree assemblies should determine which repeat architectures define exchange substrates and how these processes contribute to acrocentric chromosome evolution and rearrangement.

## Supporting information

Supplementary_discussion

Supplementary_FASTA_Sequences_Alignment

## Online Methods

### Human assemblies and K111-family classification

Human acrocentric K111-family loci were identified in haplotype-resolved Human Pangenome Reference Consortium (HPRC) assemblies using locked K111 (9,194 bp), K222 (6,254 bp) and soloLTR (968 bp) reference sequences. BLAST+ v2.12.0^15^ screening required ≥98% nucleotide identity, with initial aligned-length thresholds of ≥1,000 bp for K111/K222 and ≥800 bp for LTR queries. Final structural classification required K111 ≥8,500 bp, K222 ≥6,000 bp, intact soloLTR ≥900 bp and truncated soloLTR 800–899 bp. Competing hits were resolved by the strongest locus-specific match using single-assignment logic; duplicate, embedded LTR and redundant nested cross-hits were removed. The final population dataset comprised 46 individuals, 92 maternal and paternal haplotypes and 455 nonredundant K111-family elements (73 K111, 169 K222, 194 soloLTR and 19 truncated soloLTR). Chromosome assignments were obtained directly from HPRC assembly metadata; scaffolds lacking chromosome assignments were classified as unassigned. Reference sequences are provided in Supplementary Data 1.

### Reference sequences and validated loci

K111-family element boundaries in CHM13v2 were defined as K111, chr15:2,092,086–2,101,275; K222, chr13:5,381,823–5,388,076; soloLTR, chr14:1,698,125–1,699,093 and chr21:2,704,538–2,705,506; and truncated soloLTR, chr22:4,408,446–4,409,319. Figure-specific K111-family element boundaries were defined independently for each analysis. A 6-bp GAATTC target-site duplication was identified at the CHM13 chr15 K111 locus and the chr14 and chr21 soloLTR loci. This was treated as a candidate ancestral insertion signature and not used to infer subsequent K111-family copy-number expansion or remodeling.

### Acrocentric repeat annotation

Human acrocentric scaffold architecture was annotated using a validated 59-query library spanning ACRO, alpha-satellite HOR, BSAT, CER, HSAT1, HSAT3, SST1, rDNA and K111-family sequences. Class-specific identity and minimum-length thresholds were ACRO ≥80%/500 bp; alpha HOR ≥80%/500 bp; BSAT ≥80%/500 bp; CER ≥75%/500 bp; HSAT1 ≥85%/500 bp; HSAT3 ≥70%/500 bp; SST1 ≥80%/500 bp; rDNA ≥95%/500 bp; and K111-family ≥98%/500 bp with single assignment. Accepted hits were strand-normalized, sorted and merged according to repeat-class rules. The complete 59-query library is provided as Supplementary Data 1.

### Chimpanzee comparative genomics

Chimpanzee chromosome-resolved satellite coordinates were obtained from the supplementary mPanTro3 T2T great-ape annotation.^4^ Published annotations were simplified into the repeat classes used for scaffold comparison, and independently detected K111 loci were overlaid. Full-length chimpanzee K111 coordinates were chr14:3,024,122–3,033,302; chr15:3,156,380–3,165,559; chr17:2,865,720–2,874,901; and chr22:2,787,326–2,796,505. The six chimpanzee scaffolds retained for comparison corresponded to HSA13, HSA14, HSA15, HSA18, HSA21 and HSA22. For focused NOR-boundary analysis, repeat coordinates were used to quantify distances and block lengths across rDNA, BSAT1, BSAT2, CER and downstream repeat domains. Scaffold sequences were also queried by megablast (e-value 1 × 10^−20^) against the human and primate query libraries, and accepted hits were converted to genomic intervals for comparison of repeat order, orientation, block size and spacing.

### Phylogenetic analyses

K111 and K222 internal regions and LTRs were extracted from human pangenome assembly FASTA files using curated coordinates derived from repeat annotation and sequence-similarity searches. For LTR analyses, 5′ and 3′ LTRs from the same proviral insertion were treated as separate sequences. Extracted sequences were curated in BioEdit^16^ to verify homologous boundaries and remove artifactual extensions or poorly aligned termini.

Multiple sequence alignments were generated using MAFFT v7.505^17^ (--auto) and inspected before phylogenetic inference. The LTR phylogeny comprised 541 sequences across 1,114 alignment positions, including human K111 and K222 LTRs, soloLTR and truncated soloLTR sequences, and chimpanzee and bonobo K111 LTRs (Extended Data Fig. 3). The K111/K222 internal-sequence phylogeny comprised 249 sequences (80 K111 and 169 K222) across 6,139 alignment positions (Extended Data Fig. 2).

Maximum-likelihood trees were inferred using RAxML v8.2.12^18^ under the GTR + Γ nucleotide substitution model. Statistical support was assessed using 1,000 rapid bootstrap replicates, followed by identification of the best-scoring maximum-likelihood tree. Trees were visualized and annotated using iTOL.^19^

### Full-length K111/K222 structural analysis

Structural comparisons included 72 human full-length K111 and 168 human K222 sequences together with seven Pan K111 sequences. K222 sequences were aligned to the full-length K111 reference using sequence anchors and local sequence alignment to identify the transition into the shared K111/K222 homologous region. Boundary coordinates were compared across K222 elements. Chimpanzee and bonobo K111 sequences were examined across the corresponding interval to polarize retention or loss of the ancestral K111-like 5′ region. The short K222 sequence preceding the principal K111-collinear region was compared with individual human and Pan K111 sequences to test assignment to a distinct K111 lineage.

### K111/K222 parental-affinity and breakpoint analysis

Parental affinity was evaluated for 250 aligned K111/K222-derived sequences across positions 3,000–9,000 relative to the K111 reference. Sequence identity to K111 and K222 consensus sequences was calculated in 400-bp sliding windows containing at least 100 informative positions. Parental affinity was expressed as K222 − K111 identity, and K111-like and K222-like states were assigned using an absolute identity-difference threshold of 0.0025. Sequences containing supported transitions between states were classified as candidate mosaics.

Non-random breakpoint localization was tested using an all-site circular-shift permutation analysis that preserved the breakpoint profile while shifting its position across the analyzed sequence. A total of 1,000 permutations were performed across 240 evaluated alignment sites. Empirical *P* values were calculated from the permutation distributions and adjusted using the Benjamini–Hochberg procedure^20^ with an FDR threshold of *q* < 0.10.

Five candidate mosaic elements were examined: HG00735 PAT and HG01071 PAT, annotated as K111, and HG00290 MAT, HG00344 MAT and HG01074 MAT, annotated as K222. SimPlot^21^ analyses used 400-bp windows, 50-bp steps and 350-bp overlap. Putative breakpoint intervals were defined from transitions in parental affinity and diagnostic K111/K222 sequence markers.

Candidate recombination events were evaluated using RDP5^22^ with human K111 and K222 consensus sequences as parental references. Analyses included RDP, GENECONV, BootScan, MaxChi, Chimaera, SiScan and 3Seq. Candidate events were considered supported when detected by at least three independent algorithms and when inferred breakpoint intervals overlapped affinity transitions identified by sliding-window analysis.

### LTR parental-affinity and ancestral-state polarization

Sequence relationships among human K111 5′LTR, K111 3′LTR, K222 3′LTR and soloLTR classes were evaluated using the curated LTR alignment. The dataset comprised 191 full soloLTR sequences. One reverse-oriented HG002 PAT scaffold_44 soloLTR was excluded from sequence-level parental-affinity analyses, leaving 190 full soloLTR sequences. Consensus sequences were generated for K111 5′LTR, K111 3′LTR, K222 3′LTR and soloLTR classes, and nucleotide identity was calculated across homologous alignment positions. Individual soloLTRs were evaluated against the three proviral LTR classes. A one-switch model allowing a transition between K111 5′ and 3′LTR affinity was evaluated for a simple intraproviral 5′–3′LTR crossover.

Chimpanzee and bonobo K111 LTR sequences were used to polarize human LTR sequence changes. An ancestral Pan proxy was assigned only at alignment positions where the chimpanzee and bonobo consensuses agreed, yielding 924 Pan-polarized positions. Derived states were defined as human consensus nucleotides differing from the Pan proxy. Enrichment of positions at which K222 3′LTR and soloLTR shared the same derived state was tested by one-sided Fisher’s exact test and an exact circular-shift analysis across all 923 non-zero offsets. The empirical circular-shift *P* value was the fraction of shifted configurations with overlap greater than or equal to that observed. A stringent secondary analysis was restricted to 913 positions at which the Pan proxy and both human K111 5′ and 3′LTR consensuses carried the same nucleotide.

### Population structural-variant analysis

Population structural variation was analyzed using the CHM13v2-aligned CoLoRSdb v1.0.0^23^ pbsv/Jasmine structural-variant callset comprising 1,381 samples.

Acrocentric repeat compartments were defined from the CHM13v2 CenSat annotation for chromosomes 13, 14, 15, 21 and 22. Analyses were restricted to the acrocentric sequence preceding the annotated q-arm boundary of each chromosome. K111-family intervals were defined using the CHM13v2 loci described above.

For repeat-class comparisons, each SV was assigned by its VCF start coordinate to avoid counting long variants in multiple repeat compartments. SV density was calculated as variant start sites divided by total annotated sequence length. Rate ratios compared each repeat class with the remaining analyzed acrocentric sequence. Two-sided conditional binomial tests were used for rate comparisons. Exact 95% confidence intervals were calculated, and *P* values were adjusted across the six tested repeat classes using the Benjamini–Hochberg procedure.

Local SV burden was quantified within ±500 kb of each K111-family locus. Variants were classified by CoLoRSdb allele frequency as rare (AF <1%), intermediate (1% ≤ AF <5%) or common (AF ≥5%). Local landscapes used 110-kb windows centered on each K111-family element. Insertions and deletions were displayed according to signed SV length, with inversions shown separately where present. CenSat intervals were intersected with each window to define the surrounding repeat architecture.

To compare K111-family elements with their local repeat environment, CenSat-annotated CER sequence within 50 kb of each focal element was used as a matched background, excluding the K111-family interval. SV densities in K111-family cores and local CER were compared using a two-sided conditional binomial rate test with an exact 95% confidence interval for the rate ratio. CER sequence was further stratified by distance from the nearest element boundary (0–1, 1–2, 2–5, 5–10, 10–25 and 25–50 kb), and SV density within proximal CER (0–5 kb) was compared with distal CER (5–50 kb) using the same test.

### LTR erosion analysis

Aligned distal 3′ LTR termini were screened for recurrent terminal length changes adjacent to CER. Nested 9-bp and 12-bp distal erosion states were identified relative to the reference-length LTR. Sequence coordinates and the conserved upstream boundary were inspected to distinguish terminal erosion from generalized sequence truncation. Erosion states were summarized by individual.

### Analysis of recurrent K111-family loci and extended repeat architectures

Haplotype-resolved assemblies containing more than one K111-family element were identified from the annotations described above. For each locus, surrounding repeat annotations were extracted. CER association was assessed within 100 kb of each K111-family element. For haplotypes containing two K111-family loci, inter-locus distances were calculated from element coordinates and the intervening repeat architecture was examined.

For HG002, local CER architectures surrounding the paternal chromosome 21 soloLTR and the two soloLTR-containing regions of scaffold_44 (JAKCWS010000371.1) were compared using the lengths of the immediately flanking CER blocks. Because scaffold_44 G2 occurs in the reciprocal orientation, its flanking CER segments were orientation-normalized before comparison. Differences between architectures were calculated independently for the two CER flanks.

To determine whether sequence similarity between G1 and G2 extended beyond the focal soloLTRs, scaffold_44 was self-compared using exact 31-mers sampled at 100-bp intervals to identify extended regions of reciprocal homology. Sequence correspondence between G1 and reverse-complemented G2 was quantified using 397 exact 31-mer anchors sampled from G1 at 1-kb intervals. Significance was assessed by a scaffold-aware permutation test in which the fixed 397-kb G1 query was compared with 1,000,000 randomly positioned, matched-length windows from scaffold_44 using the same exact 31-mer procedure. The empirical P value was calculated with a +1 correction as (k + 1)/(n + 1). Identity between the focal G1 and G2 soloLTR sequences was assessed after reverse-complementing G2.

### Parental comparison of CER architectures

To test whether the HG002 paternal chromosome 21, G1 and G2 architectures resembled either parental repeat background, K111-family loci from the HG003 paternal and HG004 maternal assemblies were compared using the lengths of the CER blocks immediately flanking each element. Architectural similarity was quantified as the Manhattan distance between the two CER lengths. Because repeat-domain orientation can differ among loci, distances were calculated in both direct and reversed left–right orientations and the smaller value was retained. For each HG002 query architecture, the minimum distance to any eligible HG003 or HG004 K111-family locus was recorded.

Significance was evaluated by permutation against the parental K111-family architecture background using the same orientation-independent distance metric. The paternal chromosome 21, G1 and G2 architectures showed no significant parental preference (permutation P = 0.54335, 0.54047 and 0.53966, respectively). Parental architectural similarity alone was therefore not interpreted as evidence of parental origin or transmission.

### Trio-based analysis of K111-family transmission and structural remodeling

Six parent–offspring trios were initially evaluated: HG002/HG003/HG004, HG00609/HG00607/HG00608, HG00733/HG00731/HG00732, HG00738/HG00736/HG00737, HG00741/HG00739/HG00740 and HG00423/HG00421/HG00422. Detailed trio-resolved analyses were restricted to HG002 and HG00733 because these trios had sufficient long-read coverage (>20×) and sufficiently complete parental K111-family haplotypes for reliable offspring-to-parent comparisons. In the remaining trios, lower parental long-read coverage and/or incomplete recovery of parental K111-family loci precluded confident interpretation.

Phased offspring K111-family loci from HG002 and HG00733 were compared with K111-family loci in their corresponding parents. HG002 paternal and maternal haplotypes were evaluated against HG003 and HG004, respectively, and HG00733 paternal and maternal haplotypes against HG00731 and HG00732, respectively. Analyses incorporated K111, K222, soloLTR, truncated LTR, CER, SST1, ACRO, HSAT3 and BSAT annotations. Overlapping annotations were resolved using the priority K111/K222/soloLTR/truncated LTR > CER > SST1 > ACRO > HSAT3 > BSAT. Repeat architecture was represented in 100-bp bins and summarized across 5-kb windows.

For HG002 PAT scaffold_44 (JAKCWS010000371.1), the two soloLTR-associated domains were analyzed independently as G1 and G2 using 120,969-bp windows centered on the G1 soloLTR (1,131,709–1,132,677 bp) and G2 soloLTR (2,593,605–2,594,573 bp). This earlier HG002 assembly was retained because it contains the resolved scaffold_44 G1/G2 architecture analyzed here; the assembled organization was independently evaluated using raw ONT reads.

### Sequence and architecture similarity

Parental K111-family sequences were aligned to each offspring locus using minimap2^24^ with the Oxford Nanopore mapping preset. Sequence support was calculated within 5-kb windows from nucleotide identity and the fraction of parental sequence represented by aligned bases. Repeat-architecture similarity was calculated from informative 100-bp bins.

For each offspring-parent comparison, a composite transmission score was calculated as

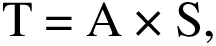

where A is repeat-architecture similarity and S is sequence support. Statistical discrimination between the best and alternative parental matches was assessed by resampling informative 5-kb windows with replacement for 10,000 bootstrap replicates and recalculating the difference in transmission score. Empirical bootstrap P values were calculated with finite-sample correction. A locus was classified as showing phase-consistent transmission support only when the best-supported parental match agreed with the independently phased parent and P < 0.05. Significant matches to the opposite parent were retained as cross-parent similarities but were not interpreted as transmission.

### Quantification of structural remodeling

Structural remodeling was evaluated independently of transmission-score ranking. For offspring loci with sufficient representatives from the expected phased parent, architecture distance was defined as

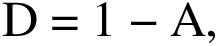

where A is repeat-architecture similarity. Remodeling was quantified as the excess offspring-parent architectural distance relative to architectural variation among representative loci from the corresponding parent:

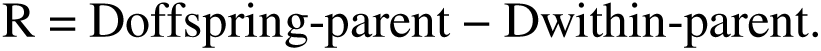

Positive R values indicate greater offspring-parent architectural divergence than expected from within-parent variation. Statistical significance was assessed by resampling informative windows with replacement for 10,000 bootstrap replicates. Loci with R > 0 and P < 0.05 were classified as showing supported remodeling. Loci lacking sufficient expected-parent representatives were not assigned a remodeling classification.

Transmission and remodeling were evaluated independently. Transmission and remodeling classifications alone were not considered sufficient evidence of de novo recombination. De novo exchange was assigned only where offspring parental-affinity transitions were supported by independent sequence-level statistical analysis and raw-molecule validation, as described below.

### Assembly-based parental sequence-affinity analysis

Local parental sequence affinity was evaluated in phased offspring K111-family loci by comparing offspring sequence independently with the paternal and maternal sequence datasets. Child loci were divided into non-overlapping 250-bp windows. For each window, nucleotide identity to the paternal and maternal datasets was calculated, and windows were classified as maternal-like, paternal-like or ambiguous from the direction and magnitude of the identity difference. Contiguous informative windows were used to identify candidate changes in parental affinity; ambiguous windows were retained in the source data but were not used to define transition direction.

For HG00733, a 475,435-bp paternal scaffold 048 child interval containing the K222-associated repeat domain was analyzed. Candidate boundaries were evaluated with a locus-wide circular-shift spatial null that preserved the observed organization of the parental-affinity signal while relocating it across eligible positions. Empirical P values were calculated with the +1 correction. These assembly-level tests were performed independently of the raw-molecule sign tests described below.

### HG002 scaffold_44 G1-G2 parental-affinity analysis

The HG002 scaffold_44 G1 and G2 regions were analyzed separately to avoid confounding parental-affinity measurements by the intervening repetitive rDNA array. The same 250-bp classification was applied independently to G1 and G2, with HG003 representing the paternal genome and HG004 the maternal genome. Ambiguous windows were excluded from the directional comparison. Differences in maternal-like and paternal-like windows between G1 and G2 were evaluated using a two-sided Fisher’s exact test. The intervening rDNA array was not used for nucleotide-resolution transition mapping.

### Raw ONT validation of trio-associated architectures

Raw Oxford Nanopore reads were analyzed independently of the haplotype assemblies to validate the HG00733 parental-affinity transitions and HG002 scaffold_44 architectures. All molecule counts were based on unique read identifiers. Unless otherwise stated, alignments were generated with minimap2 v2.31 using the map-ont preset and processed with samtools v1.24.²

### HG00733 molecule-level transition analysis

Reads from three independent HG00733 HPRC ONT libraries were recruited to the 475,435-bp paternal scaffold 048 child interval. Parental-informative sites defined by comparison of the child sequence with the paternal and maternal haplotypes were scored directly on each recruited molecule. A molecule qualified for a boundary when it contained at least four informative sites on each side. For the 229,125 boundary, the predicted direction was paternal-like to maternal-like; for the 348,375 boundary, the predicted direction was maternal-like to paternal-like. Each molecule was classified as showing a shift in the predicted direction, the opposite direction or a tie. Results were also stratified by sequencing library.

Directionality was tested using an exact two-sided sign test under the null hypothesis that a non-tied molecule was equally likely to shift in either direction. Tied molecules were excluded from the sign test. These molecule-level tests were independent of the assembly-based permutation tests.

### HG002 scaffold_44 architecture validation

The paternal HG002 scaffold_44 sequence was obtained from assembly GCA_021950905.1 (JAKCWS010000371.1). Two 120,969-bp reference windows representing G1 (scaffold coordinates 1,071,709–1,192,677) and G2 (2,533,605–2,654,573) were extracted and assigned unique sequence identifiers. Two independently generated ONT datasets were analyzed: R9.4.1 MinION/GridION reads re-basecalled with Guppy v4.2.2 and restricted to reads >50 kb, and 2023 R10.4.1 ultra-long reads. Primary R10 read sequences were extracted after excluding secondary and supplementary alignment records.

For architecture-conditioned recruitment, raw reads were recruited independently with 10-kb sequence anchors specific to G1 or G2 and aligned competitively against the combined G1 and G2 targets. Alignments were required to span at least 5 kb with MAPQ ≥20. Each molecule was assigned to the architecture with the greatest number of matching aligned bases. Because recruitment used architecture-specific anchors, these counts were treated as validation of recruited molecules and not as unbiased estimates of G1:G2 abundance.

The G2 structure was additionally evaluated by direct junction-spanning evidence. Within the G2 window, the downstream CER block ended at position 79,799 and ACRO began at position 96,118. Reads supporting the CER-bridge-ACRO interval were required to begin at or before position 69,799 and extend through at least position 106,118, providing at least 10 kb of aligned sequence on each side. Complete-path reads were required to begin at or before position 49,999, at least 10 kb upstream of the soloLTR, and extend through position 106,118, at least 10 kb into ACRO. Junction-supporting reads were primary, non-supplementary alignments with MAPQ ≥20 and were quantified independently in the R9 and R10 datasets.

### Immunofluorescence-fluorescence in situ hybridization

BJ-5Ta cells were grown to 50–70% confluency, fixed in 4% paraformaldehyde, permeabilized in PBST containing 0.2% Triton X-100 and blocked in 2% BSA/PBST. Nucleolin was detected using rabbit polyclonal anti-nucleolin antibody (Sigma-Aldrich, N2662; 15 µg/ml) for 1 h at 37 °C, followed by Alexa Fluor-conjugated secondary antibody (1:1,000) for 45 min at room temperature.

K111-family FISH probes were generated from PCR-amplified K111, K222 and soloLTR sequences.^6,7^ The 5′ K111 region was amplified using primers P1 and P4, K111/K222 3′ regions using ET1 and P2, and soloLTR sequences using P1 and P2. Probe DNA was fluorescently labeled using the ARES Alexa Fluor 488 DNA Labeling Kit (Invitrogen, Thermo Fisher Scientific, A21665) according to the manufacturer’s instructions.

Following immunostaining, cells were washed three times in PBS and treated with 100 µg/ml DNase-free RNase (New England Biolabs, T3018) for 2 h at 37 °C. Ten nanograms of fluorescent DNA probe were applied in 50% formamide, 10% dextran sulfate and 2× SSC. Probe and cellular DNA were simultaneously denatured for 2 min at 74 °C and hybridized overnight at 37 °C in a humidified dark chamber. Coverslips were washed once in 2× SSC at room temperature, followed by 2× SSC and 0.5× SSC for 5 min each at 40 °C. Nuclei were counterstained with DAPI in ProLong Gold.

Fluorescence images were acquired using an Olympus BX73 microscope equipped with a 60× objective and cellSens Dimension software under identical acquisition settings across conditions.

### Fluorescence colocalization analysis

Colocalization between K111-family probe and nucleolin fluorescence was analyzed in Fiji (ImageJ) using Coloc 2.^26^ Individual probe puncta were defined as regions of interest (ROIs), and paired probe and nucleolin fluorescence-intensity measurements were obtained from 50 probe-associated ROIs per sequence class. Pearson product-moment correlation coefficients were calculated independently for K111, K222 and soloLTR. Data were obtained from three independent experiments.

### Analysis of cells with disrupted nucleolar organization

For Extended Data Fig. 9, cytospin-prepared cells showing disrupted nuclear and nucleolar morphology were used qualitatively to assess K111-family localization following nucleolar reorganization and were not used to infer a mechanistic effect of nucleolar disruption.

### Figure preparation

Quantitative plots were generated using GraphPad Prism and Python with Matplotlib. Final figure panels were assembled without alteration of the underlying quantitative data.

### Use of artificial intelligence

OpenAI ChatGPT was used to assist with manuscript editing, language refinement, code development and evaluation of analytical and statistical approaches. All analyses, statistical tests, results, interpretations and manuscript content were independently reviewed and verified by the authors. The authors take full responsibility for the accuracy and integrity of the work.

### Statistics and reproducibility

Statistical tests, sample sizes and resampling replicate numbers are specified in the relevant Online Methods subsections and figure legends. All tests were two-sided unless explicitly identified as one-sided. Exact and empirical P values are reported without rounding to zero. Empirical permutation and bootstrap P values used finite-sample correction, (k + 1)/(n + 1). Multiple-testing correction used the Benjamini-Hochberg procedure with the false-discovery-rate threshold stated for the corresponding analysis. No statistical method was used to predetermine sample size, and no data were excluded unless the exclusion criterion is described in the relevant method.

Population composition was evaluated by PERMANOVA^26^ using Bray-Curtis distances and 9,999 permutations, followed by class-specific Kruskal-Wallis tests with Benjamini-Hochberg correction. Chromosome-level distributions of K111-family structural classes were compared using a chi-square test of independence. Paired maternal-paternal copy-number comparisons used two-sided Wilcoxon signed-rank tests. Associations among K111-family class counts used Pearson product-moment correlation.

No inferential comparison was made between HG002 R9 and R10 read counts or between architecture-conditioned G1 and G2 recruitment counts. No inferential comparison was performed among the three IF–FISH correlation coefficients.

## Data availability

The T2T-CHM13 v2.0 human reference genome used in this study is available from NCBI under GenBank assembly accession GCA_009914755.4 (BioProject PRJNA559484). Haplotype-resolved human genome assemblies were obtained from Human Pangenome Reference Consortium (HPRC) public resources. The HG00733 maternal and paternal assemblies used for trio analyses are available under accessions GCA_018506975.2 and GCA_018506955.2, respectively. The specific assemblies and contigs used in population and locus-level analyses are identified in the Supplementary Tables and Supplementary Data.

For the HG002 scaffold_44 analysis, we additionally used the earlier HG002/NA24385 paternal assembly GCA_021950905.1, specifically scaffold_44 (JAKCWS010000371.1). This scaffold was retained because the K111-family locus analyzed here is represented in this earlier assembly but is not equivalently represented in the later HG002 assembly release used for the other analyses.

Comparative great-ape T2T genome assemblies for Pan troglodytes, Pan paniscus, Gorilla gorilla, Pongo abelii, Pongo pygmaeus and Symphalangus syndactylus were obtained from NCBI genome resources. The sequence and assembly accessions used for these comparative analyses are provided in Supplementary Table 1.

Structural-variant analyses used CoLoRSdb v1.0.0 on the CHM13v2 reference (CoLoRSdb.CHM13.v1.0.0.pbsv.jasmine.vcf.gz), comprising 1,381 samples with data.

No new sequencing data were generated in this study. Raw Oxford Nanopore sequencing data used for parental comparisons and molecule-level validation were obtained from previously generated public datasets. Parental sequence analyses used HG003 (ERR17533525), HG004 (ERR17533528), HG00731 (ERR12954391) and HG00732 (ERR13491767) ONT datasets. HG00733 offspring validation used three independent ONT libraries obtained through the HPRC PLUS public data resource (HG00733_1, HG00733_2 and HG00733_3, corresponding to Libraries 1, 2 and 3, respectively, in Fig. 7 and Extended Data Fig. 8) and associated with BioProject PRJNA731524. HG002 scaffold_44 validation used publicly available Genome in a Bottle/Human Pangenome Project ONT data for HG002/NA24385, including MinION/GridION R9.4.1 reads re-basecalled with Guppy v4.2.2 and an independently generated R10.4.1 ultra-long dataset.

The K111-family reference sequences and repeat-query library used for sequence identification and annotation are provided in Supplementary Data 1. Sequence datasets underlying the phylogenetic and recombination analyses are provided in Supplementary Data 2 and Supplementary Data 3. Numerical data underlying all main and Extended Data figures are provided as Source Data files. Raw IF–FISH microscopy images underlying Fig. 8 and Extended Data Fig. 9 are available from the corresponding author upon reasonable request. All other data supporting the findings of this study are available within the Article and its Supplementary Information.

## Code availability

Custom scripts used for repeat classification, remodeling and transmission scoring, permutation analyses, trio ancestry analysis and Oxford Nanopore read-level validation are available through Zenodo at doi:10.5281/zenodo.22712456. Publicly available software, software versions, analysis parameters and statistical procedures are described in the Methods. Where original historical scripts were not retained, reconstructed implementations validated against archived source data are provided and clearly labeled as reconstructed.

## Acknowledgements

The authors acknowledge the University of Alabama at Birmingham IT-Research Computing group for high-performance computing support and computational resources provided by the Cheaha compute cluster.

## Author contributions

M.W. performed experimental studies, including IF–FISH, carried out comparative primate analyses, analyzed data and contributed to writing the paper. A.D. prepared genomic scaffolds and performed IF–FISH experiments. H.O. performed phylogenetic analyses. A.I. performed IF–FISH experiments. R.C.-G. conceived and designed the study, supervised the project, performed computational analyses, interpreted the data, acquired funding and wrote the paper. All authors reviewed and approved the manuscript.

## Additional information

Supplementary Information is available for this paper.

Correspondence and requests for materials should be addressed to Rafael Contreras-Galindo.

---

**Extended Data Fig. 1 | K111-family composition across human populations. a,** Heatmap showing the fraction of K111-family calls represented by K111, K222, soloLTR and truncated soloLTR in each of 46 individuals, grouped by population. HG002 (AJ) is shown for completeness but was excluded from population-level inferential testing because it is represented by a single individual. **b,** Mean per-individual fraction of each K111-family class across the five 1000 Genomes populations, with 95% bootstrap confidence intervals. Global differences in four-class composition were tested by PERMANOVA using Bray-Curtis distances and 9,999 permutations (P = 0.847). Class-specific differences were tested by Kruskal-Wallis tests with Benjamini-Hochberg correction across the four K111-family classes (K111, q = 0.995; K222, q = 0.995; soloLTR, q = 0.995; truncated soloLTR, q = 0.191). No population-level comparison remained significant after multiple-testing correction. Source data are provided as a Source Data file.

**Extended Data Fig. 2 | Phylogenetic diversification of K111 and K222 internal sequences.** Phylogenetic analysis of 249 aligned K111/K222 internal sequences, comprising 80 K111 and 169 K222 sequences across 6,139 alignment positions. Human K111 sequences resolve into nine labeled phylogenetic groups (K111a–i), whereas the more numerous K222 sequences are distributed across 19 groups (K222a–s). A mixed K111/K222 lineage is also present (K111/K222a). Chimpanzee and bonobo K111 sequences provide evolutionary context for the human lineages. Branch colors distinguish phylogenetic lineages sharing supported common ancestral nodes, and black-circle size indicates bootstrap support. Source data are provided as a Source Data file.

**Extended Data Fig. 3 | Phylogenetic diversification of K111-family LTR sequences.** Phylogenetic analysis of 541 LTR sequences comprising human K111 and K222 5′ and 3′ LTRs, soloLTRs and truncated soloLTRs, together with chimpanzee and bonobo K111 LTRs. Branch colors distinguish phylogenetic lineages sharing common ancestral nodes, and black-circle size indicates bootstrap support. Human soloLTRs resolve into eight phylogenetic branches (soloLTRa–h), demonstrating substantial sequence diversity rather than formation of a single soloLTR lineage. K222 3′LTRs similarly resolve into seven branches (K222 3′LTRa–g), while K111 5′ and 3′LTR sequences occupy multiple phylogenetic branches. Chimpanzee and bonobo 5′LTRs cluster with a subset of human K111 5′LTR lineages, whereas primate 3′LTRs occupy distinct branches. Source data are provided as a Source Data file.

**Extended Data Fig. 4 | Structural-variant landscapes across K111-family loci.** Structural variants within 110-kb windows centered on the five CHM13v2 K111-family loci: **a**, chr13 K222; **b**, chr14 soloLTR; **c**, chr15 K111; **d**, chr21 soloLTR; and **e**, chr22 truncated soloLTR. CoLoRSdb v1.0.0 structural variants (1,381 samples with data) are plotted by genomic position and variant size, with insertions above and deletions below the baseline. CenSat-annotated CER sequence is shown in light blue and the focal K111-family element in red. The respective windows contained 90, 138, 168, 110 and 26 structural variants, whereas only 3, 0, 1, 0 and 0 variant start sites occurred within the focal elements, respectively. Source data are provided as a Source Data file.

**Extended Data Fig. 5 | K111-family sequences are depleted for structural variation relative to surrounding CER. a**, Structural-variant density within the five K111-family elements compared with CenSat-annotated CER sequence within 50 kb of the elements. K111-family cores showed a 6.5-fold lower SV density than local CER (rate ratio (RR) = 0.155, 95% CI = 0.042–0.399, P = 5.66 × 10^−7^). **b**, CER SV density as a function of distance from the nearest K111-family element boundary. SV density within 0–5 kb did not differ significantly from CER 5–50 kb away (RR = 0.90, P = 0.49), providing no evidence for increased structural variation specifically at K111-family boundaries. SVs were assigned by their VCF start coordinate. Two-sided conditional binomial rate tests were performed using SV counts and analyzed sequence lengths. Source data are provided as a Source Data file.

**Extended Data Fig. 6 | Paired K111-family loci show variable spacing and local CER organization. a,** Genomic separation between paired K111-family loci on six haplotype-resolved scaffolds. Bars show the distance between the two elements on each haplotype, with K111-family classes indicated. Five pairs are separated by 1.46–3.30 Mb, whereas the two K222 elements on HG00235 PAT are separated by 24.3 kb. **b,** Total CER sequence within ±100 kb of each K111-family element for the two loci on each haplotype. CER abundance differs between paired loci on five of the six haplotypes. **c,** Distribution of CER sequence on the left and right sides of each of the 12 K111-family elements within ±100 kb. Stacked bars show left-and right-side CER coverage, revealing substantial variation in both CER abundance and sidedness among elements. Element number and K111-family class are indicated below each bar. Source data are provided as a Source Data file.

**Extended Data Fig. 7 | Parental sequence affinity and raw-read validation of HG002 scaffold_44. a,** Parental affinity across the reciprocal G1 and G2 domains flanking the rDNA array. Among non-ambiguous 250-bp windows, G1 contained 24 maternal-like and four paternal-like windows, whereas G2 contained one maternal-like and 49 paternal-like windows (two-sided Fisher’s exact test, P = 6.01 × 10^−15^). **b,** G2 CER–soloLTR–CER–intervening sequence–ACRO architecture and raw-read validation intervals. **c,** Alignments of four R9.4.1/Guppy v4.2.2 and six R10.4.1/2023 ultra-long reads satisfying the complete G2-path criterion. **d,** Unique junction-supporting molecules. Forty-four R9 and 70 R10 molecules spanned the CER-bridge-ACRO interval; four R9 and six R10 molecules traversed the complete G2 path. **e,** Architecture-conditioned competitive alignments. All 774 evaluable G1-anchor-recruited molecules favored G1; among 619 evaluable G2-anchor-recruited molecules, 363 favored G2 and 256 favored G1. Recruitment counts are descriptive and do not estimate G1:G2 abundance. Source data are provided as a Source Data file.

**Extended Data Fig. 8 | Independent raw-molecule validation of HG00733 de novo maternal–paternal sequence exchange. a,** Per-library molecule support for the paternal-like to maternal-like transition at position 229,125. Qualified molecules contained at least four parental-informative sites on each side. Predicted-direction support was observed for 12 of 12 molecules in library 1, 21 of 23 in library 2 and 18 of 18 in library 3; pooled support was 51 of 53 molecules (exact two-sided sign test, P = 3.18 × 10^−13^). **b,** Per-library raw-molecule support for the maternal-like to paternal-like ancestry transition at position 348,375. Predicted-direction support was observed for 24 of 27, 26 of 28 and 27 of 30 qualified molecules in libraries 1–3, respectively. Pooled support was 77 of 84 non-tied molecules (exact two-sided sign test, P = 5.14 × 10^−16^); one tied molecule was excluded. Assembly-based circular-shift permutation P values were 0.000600 and 0.019398, respectively. Source data are provided as a Source Data file.

**Extended Data Fig. 9 | K111-derived sequences remain associated with nucleolin in cells with disrupted nucleolar organization.** Representative IF–FISH images showing K111 (top), K222 (middle) and soloLTR (bottom) localization in BJ-5Ta cells exhibiting disrupted nuclear and nucleolar organization following cell detachment and cytospin preparation. K111-family FISH signals are shown in green, nucleolin in red and DNA in blue (DAPI). Insets show representative nuclei and arrowheads indicate K111-family signals associated with redistributed nucleolin-positive domains. Scale bar, 2 µm. Images are representative of three independent experiments. Source data are provided as a Source Data file.

