## Supplementary figures and images for "Human-specific remodeling of an endogenous retrovirus shapes structural diversity at acrocentric nucleolar organizer regions"

### Extended Data Figure 1

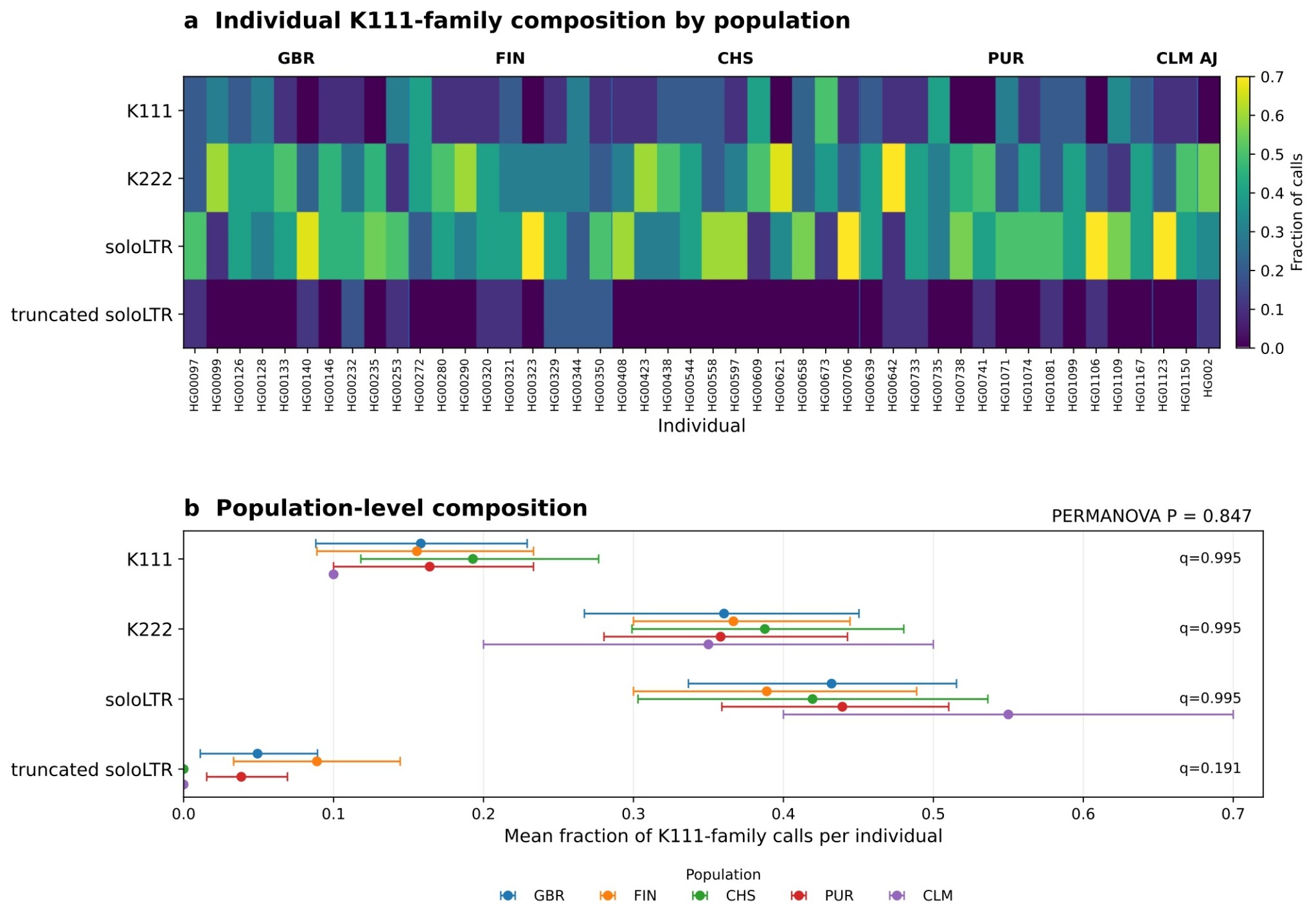

### Extended Data Figure 2

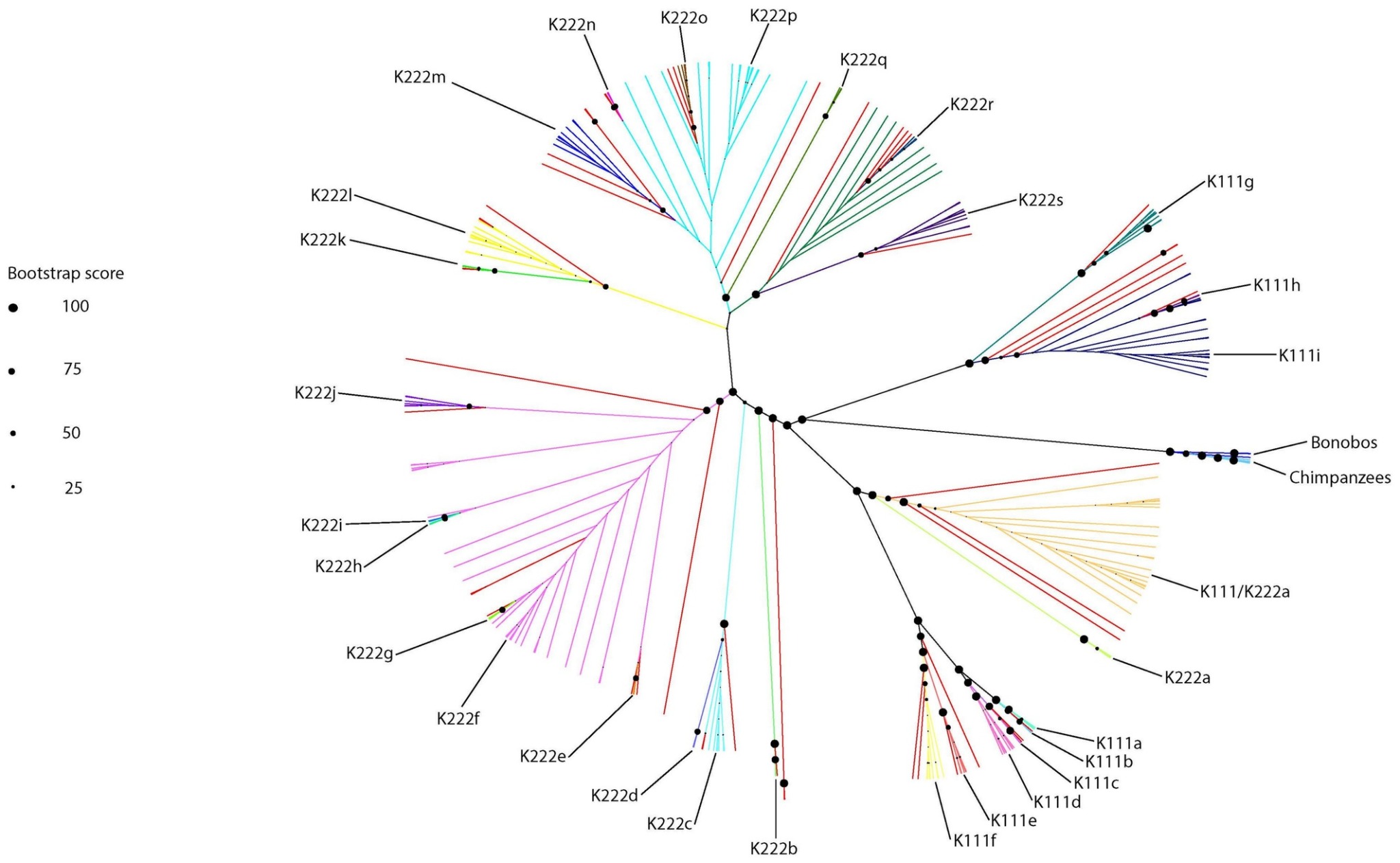

### Extended Data Figure 3

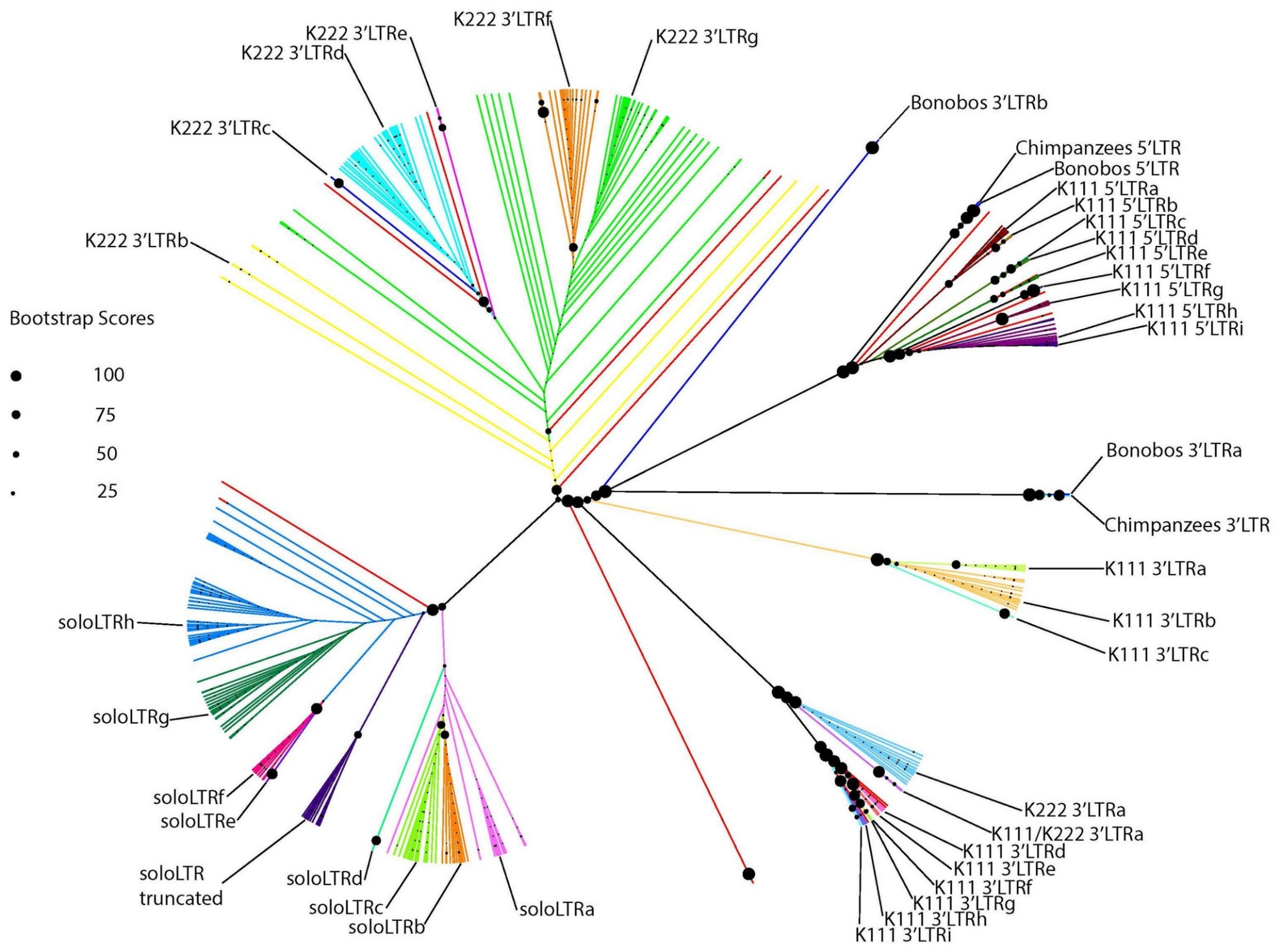

### Extended Data Figure 4

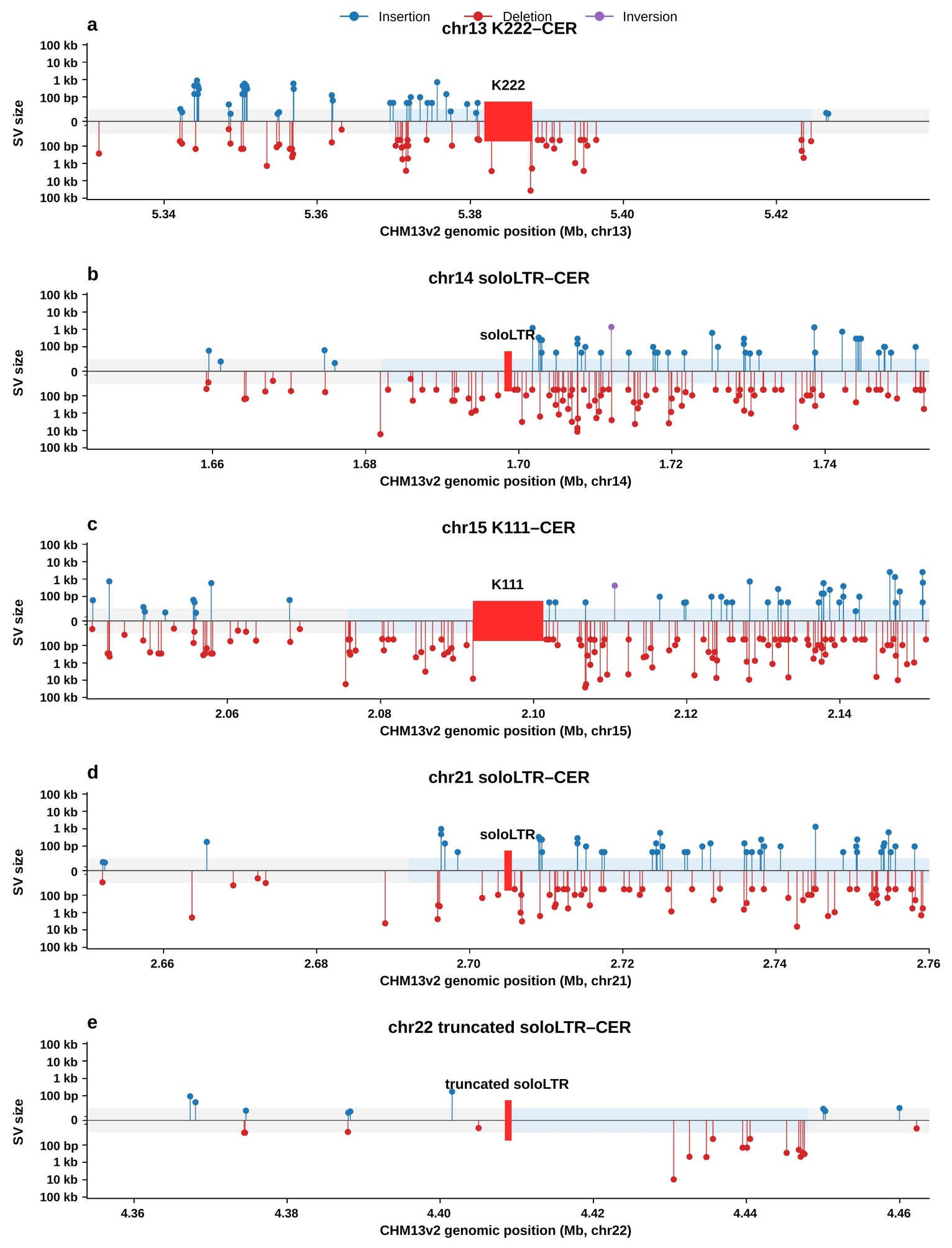

### Extended Data Figure 5

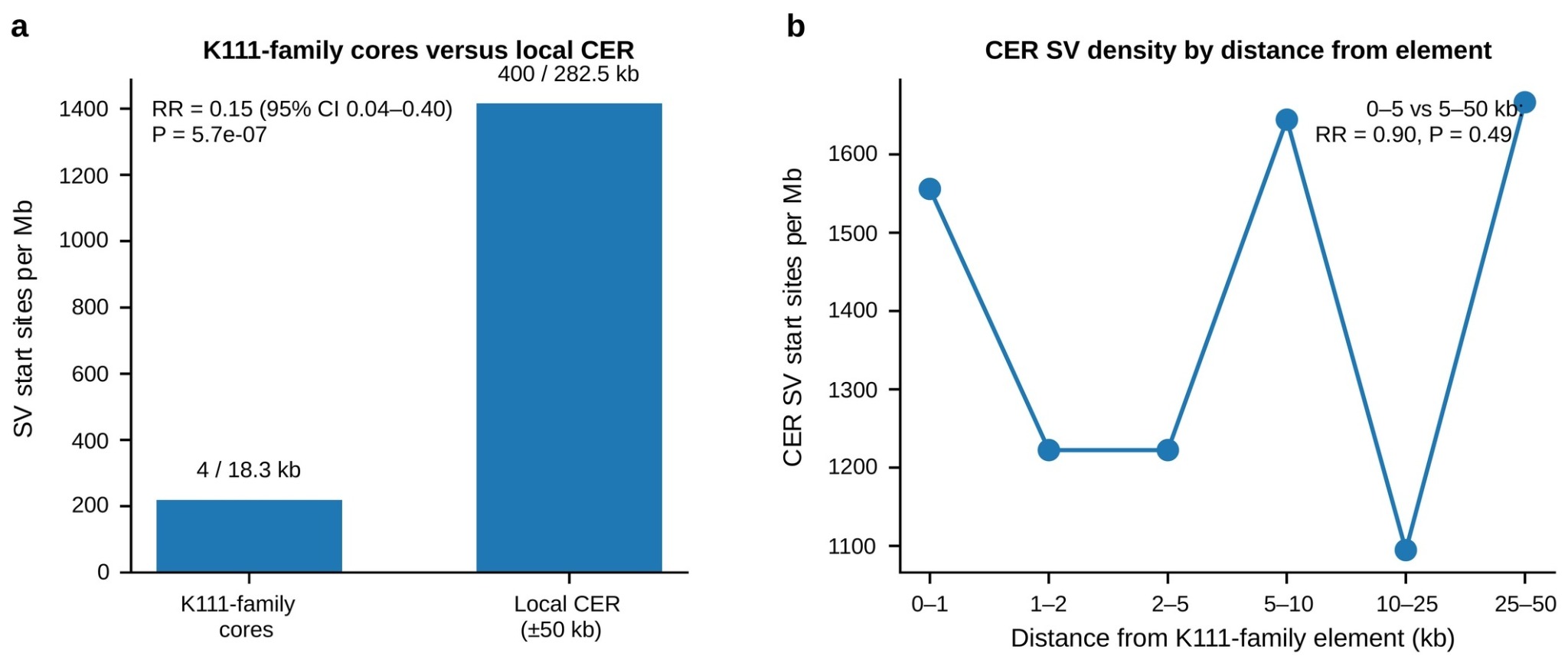

### Extended Data Figure 6

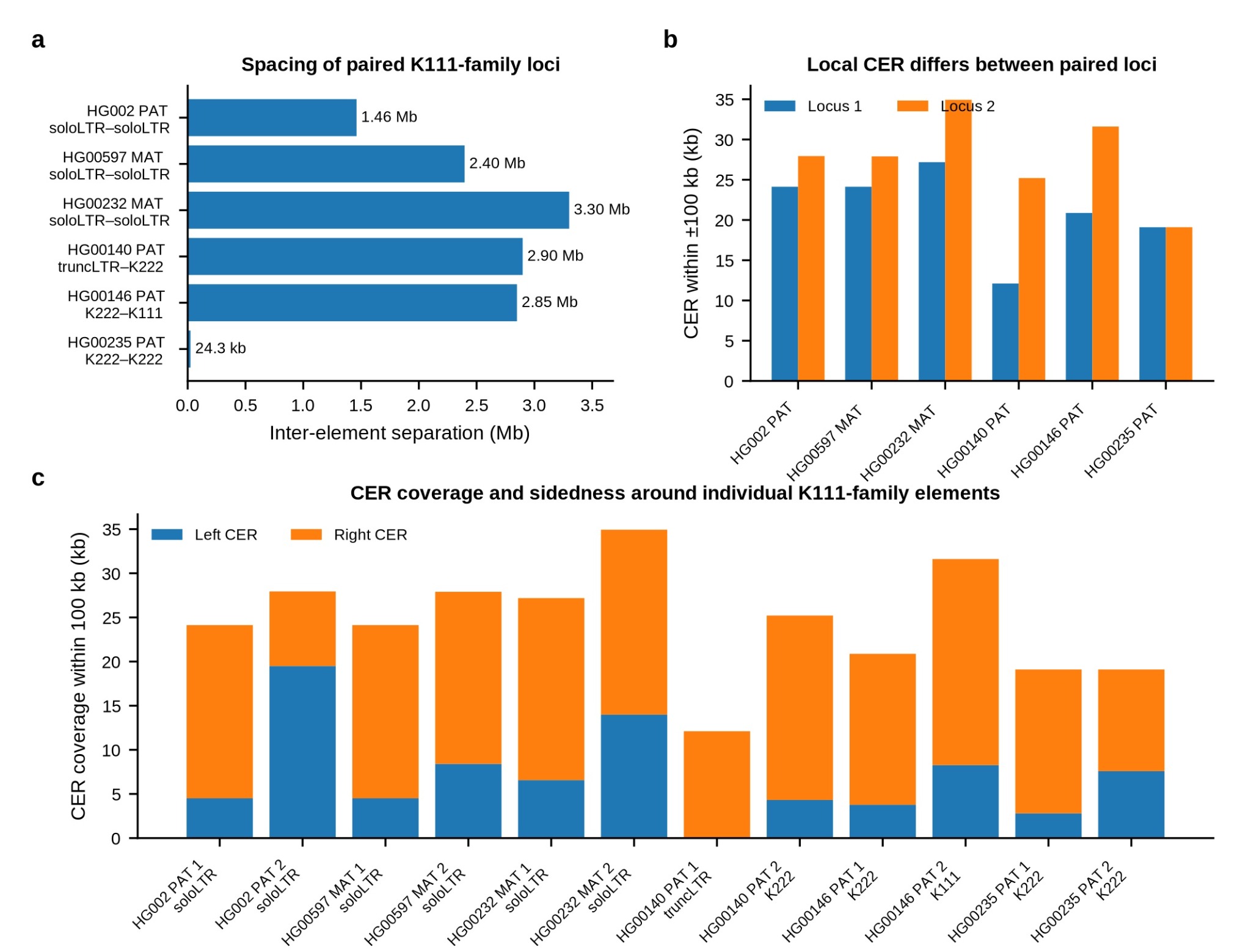

### Extended Data Figure 7

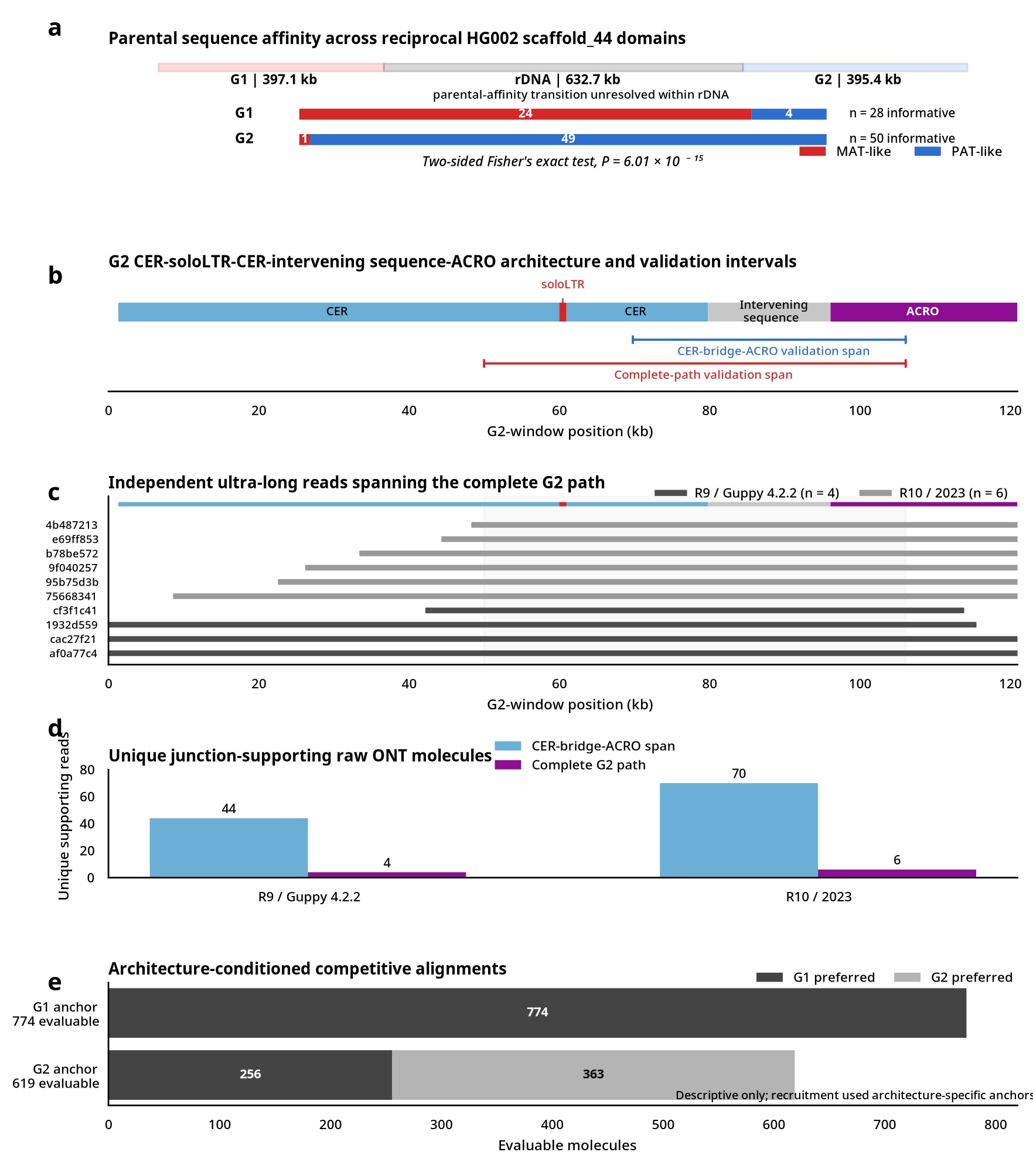

### Extended Data Figure 8

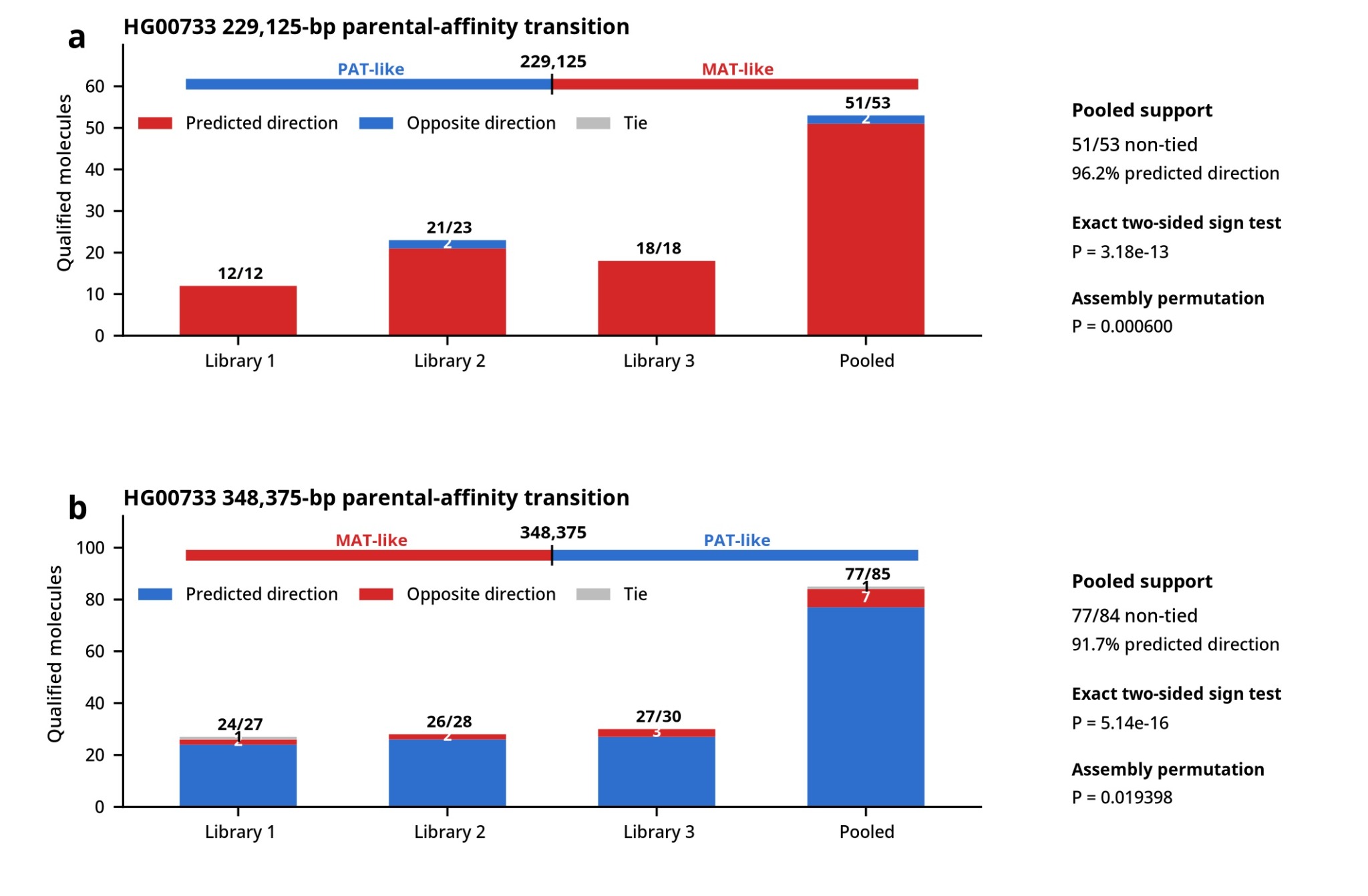

### Extended Data Figure 9

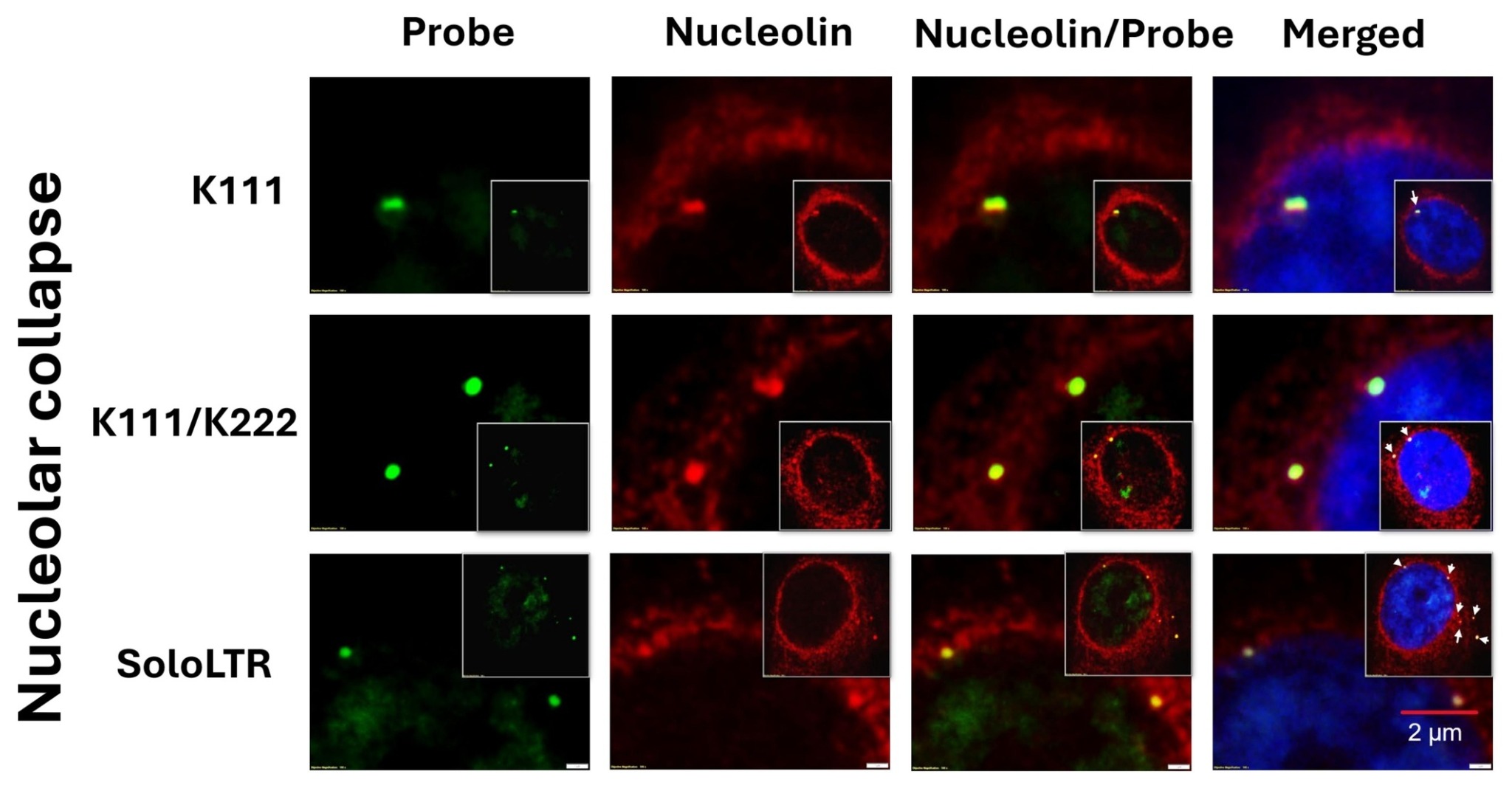
