## Supplementary_discussion for "Human-specific remodeling of an endogenous retrovirus shapes structural diversity at acrocentric nucleolar organizer regions"

**Supplementary Discussion 1 | Acrocentric architecture, nucleolar organization and chromosome rearrangement**

The combination of extensive sequence homology and shared nuclear localization provides a potential framework for understanding why particular acrocentric domains undergo exchange. Population-scale analyses have identified pseudo-homologous regions shared among heterologous acrocentrics, supporting recurrent sequence exchange despite suppression of conventional allelic recombination across much of their short arms.^3,5^ The K111-family loci analyzed here extend this view by marking homologous sequence relationships embedded within repeat architectures that can differ substantially in CER abundance, orientation and neighboring satellite composition. Thus, homology relevant to acrocentric exchange need not correspond to invariant repeat architecture.

The HG002 scaffold_44 illustrates this distinction. Its reciprocal G1 and G2 domains share sequence correspondence across hundreds of kilobases and contain nearly identical soloLTRs, yet differ in local repeat composition and parental affinity. These observations are compatible with remodeling of extended homologous domains rather than duplication and stable inheritance of small K111–CER units. Changes in CER abundance or orientation could alter the length and organization of homologous substrates without eliminating sequence relationships across the broader domain. Our data do not establish the molecular mechanism generating these architectures, and scaffold_44 should therefore not be interpreted as evidence for a specific inversion or crossover event.

Spatial organization may provide an additional constraint. NORs from multiple acrocentric chromosomes associate within nucleoli, bringing otherwise heterologous chromosomes into a shared nuclear compartment.^3,9,11,12^ The association of K111, K222 and soloLTR signals with nucleolin-positive domains therefore places structurally distinct K111-family loci in an environment where different NOR-bearing chromosomes can approach one another. This observation is compatible with a model in which nuclear proximity increases opportunities for interactions between homologous repeat domains, but it does not demonstrate that nucleolar association initiates recombination or that K111-family sequences themselves constitute recombination substrates.

These features may also be relevant to acrocentric chromosome rearrangements. Complete assemblies of human Robertsonian chromosomes recently identified recurrent breakpoints within SST1 arrays embedded in larger shared homology domains and supported a model in which sequence homology, acrocentric spatial proximity and meiotic recombination contribute to formation of common Robertsonian chromosomes.^14^ The K111-family domains characterized here are not mapped Robertsonian-translocation breakpoints, and our data do not establish a role for K111, K222 or CER in Robertsonian chromosome formation. Rather, they identify an additional class of sequence-resolved markers showing that NOR-associated acrocentric domains can combine extensive interchromosomal homology, local repeat remodeling and shared nucleolar association.

A broader model is therefore that acrocentric exchange depends on the intersection of at least three properties: sufficient sequence homology to provide an exchange substrate, local repeat architecture that permits or constrains alignment, and spatial organization that brings heterologous acrocentrics into proximity. The relative contribution of each remains unresolved. Complete multigenerational assemblies and sequence-resolved rearrangement breakpoints should allow these factors to be tested independently and determine whether the architectures identified through K111-family loci overlap preferential substrates for physiological inter-acrocentric exchange or pathological chromosome rearrangement.
